# Specialized mesenteric lymphatic capillaries by-pass the mesenteric lymph node chain to transport peritoneal antigens directly into mediastinal lymph nodes

**DOI:** 10.1101/2023.07.11.548304

**Authors:** Esther Redder, Nils Kirschnick, Shentong Fang, Michael Kuhlmann, Alejandra González-Loyola, Tania Wyss, Martin Stehling, Ralf H. Adams, Tatiana V. Petrova, Kari Alitalo, Antal Rot, Friedemann Kiefer

**Affiliations:** University of Münster, European Institute for Molecular Imaging (EIMI), Münster, Germany; Mammalian Cell Signalling Laboratory, Max Planck Institute for Molecular Biomedicine, Münster, Germany; Wihuri Research Institute, Biomedicum Helsinki, Helsinki, Finland; Translational Cancer Medicine Program, Faculty of Medicine, University of Helsinki, Finland; Department of Oncology, University of Lausanne and CHUV, Epalinges, Switzerland; Ludwig Institute for Cancer Research Lausanne, Epalinges, Switzerland; Flow Cytometry Unit, Max Planck Institute for Molecular Biomedicine, Münster, Germany; Max Planck Institute for Molecular Biomedicine, Department of Tissue Morphogenesis, University of Münster, Faculty of Medicine, Münster, Germany; Centre for Microvascular Research, William Harvey Research Institute, Barts and The London School of Medicine and Dentistry, Queen Mary University of London; Centre for Inflammation and Therapeutic Innovation, Queen Mary University London, EC1M 6BQ, London, United Kingdom

**Keywords:** Lymphangiogenesis, mesentery, Vegfc, lymphatic capillary, peritoneal immunity

## Abstract

Lymphatic vessels (LVs) are indispensable for tissue fluid homeostasis and immune cell trafficking. The network of LVs that channel fluids from the gut into mesenteric lymph nodes (MLN) has been recognized as the sole lymphatic system in the mesentery. Here we describe an alternative, functionally autonomous set of capillary mesenteric LVs (capMLVs) that by-pass the MLNs and drain directly into mediastinal LNs. CapMLVs develop perinatally from valves of collective mesenteric lymphatic vessels (colMLVs) in response to arterial endothelial cell-derived VEGF-C. Once extended, capMLVs detach from colMLVs to form an independent elongated network comprised of LYVE1+, CCL21+ endothelial cells. Avascular areas of the mesentery juxtaposed to capMLVs contain cell islets that express ACKR4. This CCL21-scavenging atypical receptor facilitates the migration of mesenteric phagocytes into capMLVs to be channeled directly into mediastinal LNs. This allows peritoneum-derived ominous antigens to be processed separately from alimentary antigens.

## Introduction

The body’s largest serous visceral cavity, the peritoneal cavity, is formed by parietal and visceral mesothelium, covering the abdominal wall, diaphragm and intraperitoneal organs. The pouch formed by mesothelium and connective tissue, the mesentery, suspends the contractile intestine within the abdominal cavity, prevents direct adhesive contacts with the parietal wall and maintains systemic connectivity through regularly organized nerves, blood and lymphatic vessels (LVs) (Isaza-Restrepo et al., 2018). Besides preventing tissue adhesion, the mesothelium controls fluid, solute and cell transport from the peritoneal cavity and generates a physiological barrier that contributes to the induction and regulation of immune responses (Coffey and O’Leary, 2016; Liu et al., 2020).

Ingested and digested in the upper gastrointestinal tract nutrients are absorbed in the gut into blood vessels to enter the liver via the portal vein, while fat-soluble dietary components, including food antigens, access intestinal lacteals and reach the thoracic duct after filtration and scrutiny in multiple mesenteric lymph nodes (MLN) (Bernier-Latmani and Petrova, 2017). Lacteals are highly permeable lymph capillaries comprised of oak-leaf shaped lymphatic endothelial cells (LECs) that have discontinuous “button” junctions and express the hyaluronan receptor LYVE1 (Baluk et al., 2007; Bernier-Latmani et al., 2015). Lacteals are closely associated with surrounding smooth muscle cells (SMC) that produce the major lymphangiogenic growth factor vascular endothelial growth factor (VEGF)-C (Karkkainen et al., 2004; Nurmi et al., 2015; Paavonen Karri 2002). Their contractions propel the newly formed lymph into the submucosal LV network (Choe et al., 2015). Further drainage via collecting mesenteric LVs (colMLVs) and mesenteric lymph nodes (MLNs) delivers the lymph into a dilated sac, called cysterna chylii, at the lower end of the thoracic duct. The ColMLVs feature spindle-shaped LECs with continuous “zipper” junctions (Baluk et al., 2007). In contrast to dietary components, peritoneal fluid and its particulate components are drained via stomata, pore-like openings in the mesothelium that have been proposed to open directly to mesenteric LVs (Isaza-Restrepo et al., 2018; Jackson-Jones and Benezech, 2020). Highly vascularized, secondary lymphoid structures, termed fat-associated lymphoid clusters (FALCs), underlie the mesothelium. FALCs are rich in stomata and localize proximal to the major mesenterial arterial, venous and lymphatic vessel bundles. They contribute to peritoneal immune surveillance, particularly through innate B-cells (Cécile Bénézech, 2015; Cruz-Migoni and Caamano, 2016). However, the exact microanatomical relations of stomata, FALCs and MLVs remains unclear.

LVs channel immune cells and antigen into lymph nodes (LNs) to initiate adaptive immune responses or to promote tolerance (Hampton and Chtanova, 2019). Phagocytes presenting food or microbial antigens and remnants of apoptotic epithelial cells are constitutively carried from the intestinal epithelium towards the MLNs, which were identified as key sites of oral tolerance induction (Huang et al., 2000; Kraus et al., 2005; Worbs et al., 2006). Importantly, MLNs preserve ignorance to commensal gut microbiota. The chain of MLNs is compartmentalized, as anatomically and physiologically distinct intestinal parts drain to segregated MLNs (Esterhazy et al., 2019; Houston et al., 2016). In contrast, the precise drainage route for peritoneal antigens remains unknown.

Development of MLVs initiates with LECs forming a primordial lymph sac at the root of the mesentery (Sabin, 1902; van der Putte, 1975). Sprouts from the lymph sac and distantly arising LEC clusters, derived from hemogenic endothelium, merge around the mesenteric blood vessels and form primitive LVs that grow towards and invade the intestinal mucosa at embryonic day of development (E) 14.5 to 15.5, coinciding with the formation of intestinal villi (Kim et al., 2007; Mahadevan et al., 2014; Stanczuk et al., 2015). The lymphatic vascular plexus remodels into mature mesenteric collectors until birth, and the lacteal growth into intestinal villi occurs around E17.5 in preparation of postnatal lipid absorption (Kim et al., 2007; Norrmen et al., 2009; Sabine et al., 2012). LV development indispensably relies on the VEGF-C - VEGFR-3 signalling axis (Tammela and Alitalo, 2010). VEGF-C is required for perinatal lymphangiogenesis, but also for LV maintenance and regulation of intestinal LV growth in adults (Nurmi et al., 2015).

Here we describe the hitherto not recognized network of mouse mesenteric capillary LVs (capMLVs), which extend from the intestinal wall through the entire mesentery and consist exclusively of oak-leaf shaped, LYVE1+, CCL21+ LECs, connected by button junctions. The development of capMLV starts at E18.5 from the valve territories of lymphatic collectors, resulting in an independent network by P28. The number of capMLVs is strongly diminished in *Vegfc*^+/LacZ^ mice, indicating an essential role of VEGF-C - VEGFR3 pathway in their expansion. We identified the arterial endothelium as a major source of VEGF-C for capMLV development, whereas collecting LVs were largely unaffected by arterial endothelial cell (EC)- specific *Vegfc* deletion. The capMLVs do not drain to MLNs, but channel cells from the peritoneal cavity into the mediastinal LNs. Avascular regions of the mesentery localizing in juxtaposition to capMLVs express ACKR4, an atypical receptor known to efficiently scavenge CCL21. Phagocyte immigration from the mesentery was impaired in *Ackr4*-deficient mice suggesting its involvement in shaping functional CCL21 patterns around capMLVs and in agreement with previously reported scavenging of CCR7 ligands like CCL21 by Ackr4 (Bryce et al., 2016; Forster et al., 2008; Friess et al., 2022; Ulvmar et al., 2014).

Taken together, we describe a novel mesenteric lymph vessel network that contributes to peritoneal immune surveillance by trafficking antigen presenting phagocytes to mediastinal LNs. Such dichotomy of lymphatic vascular networks in the mesentery allows for a spatial and functional separation of dietary and peritoneal immunity by channelling cells and antigens into alternative, functionally divergent mesenteric are mediastinal LNs.

## Results

### Continuous lymphatic capillaries extend along the blood vascular bundles throughout the entire mesentery

Lymphatic capillary vessels (capLVs) are comprised of LYVE1+ LECs connected by discontinuous button junctions (Baluk et al., 2007). In mice, the detailed anatomy of the mesenteric capMLVs has not been described so far. To identify capMLVs, we immunostained adult mesenteries for LYVE1, which revealed an interconnected capLV network, extending between the intestinal wall and the MLNs (Fig. 1 A, B). capMLVs formed distinct vessels that were fully segregated from colMLVs (Fig. 1 C) and closely aligned with the mesenteric arteries, following their bifurcations (Fig. 1 A - C). In contrast to other tissues including dermis, meninges and intestinal serosa, capMLVs were particularly elongated and only rarely branched (Fig. 1 A, B). In accord with their capillary nature, capMLVs displayed typical blind ends (Fig. 1B arrowheads). Furthermore, they lacked intraluminal valves and smooth muscle cells, indicating complete absence of collecting LV features (Fig. 1 C, D). capMLVs were not connected to FALCs (Fig. 1A, B, Fig. S1 A). They were comprised of oak-leaf shaped, LYVE1+ LECs with discontinuous button junctions (Fig. 1 E, F). We verified our immunostaining results by FACS (Fig. S1 B, C) and single cell RNA sequencing (Fig. 1 G-I) of mouse mesenteries (Gonzalez-Loyola et al., 2021). Transcriptomic analysis identified 15 mesenteric cell populations, one of which contained mesenteric capillary LECs (capMLECs, Fig. 1 G). capMLECs comprised approx. 60% of the total MLECs (Fig. S1 D), expressed VE-cadherin (Fig. S1 D), PROX1 (Fig. 1 H-I, Fig. S1 D) and the CCL21 chemokine (Fig. 1I), but not the atypical chemokine receptor ACKR4 (Fig. 1 F, H) (Friess et al., 2022). Hence, the LYVE1+, CCL21+ MLEC fraction represented bona fide capLECs, positioned independently of the stomata, widely over the entire mesothelial surface of the mesentery, including the large avascular areas between the radial, fat-associated vascular bundles extending from the central MLNs to the intestinal wall (Fig. S1 E, F - F’’).

**Figure 1.**
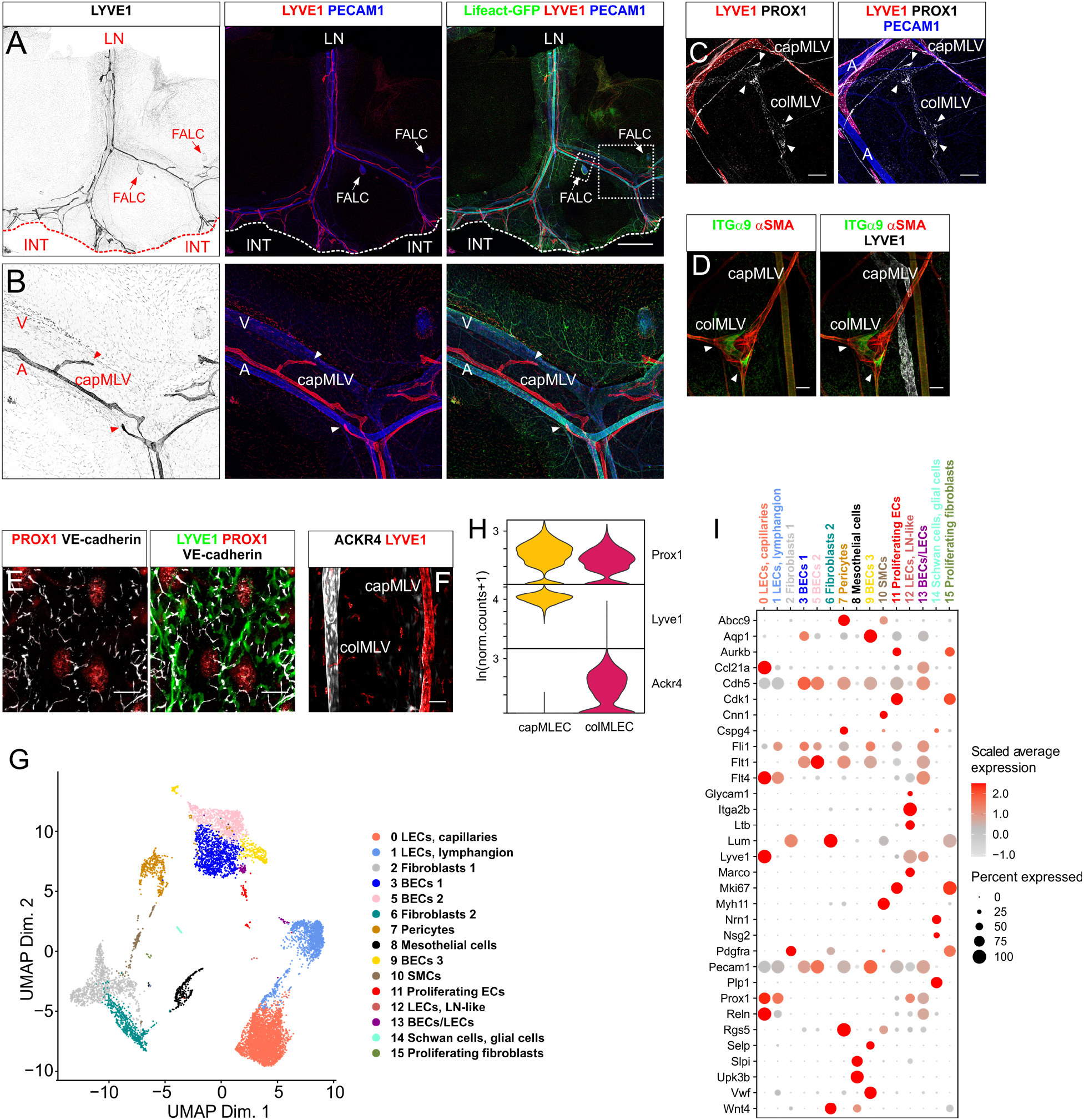
Elongated lymphatic capillaries extend between the intestinal wall and the mesenteric root. **A** Representative confocal tile scan (maximum intensity projection, MIP) of a mesenteric wholemount preparation from a 20 wks old, mouse (lifeact-GFP transgenic), immunostained for the indicated antigens. (LN = position of the removed mesenteric lymph node chain, FALC = fat-associated lymphoid cluster, INT = position of the removed intestine). White boxed areas are magnified in (B) and (Fig. S1 E). The white or red dotted line marks the attachment site of the mesentery to the small intestine. Scale bar = 2000 µm. **B** Magnified view of the mesenteric vascular network. Arrowheads indicate blind ending mesenteric lymphatic capillaries (capMLV = capillary mesenteric lymphatic vessel, A = artery, V = vein). **C** Confocal tile scan (MIP) showing distinct localization of capMLV and colMLV (collecting mesenteric lymphatic vessels). White arrowheads point to intraluminal valves in colMLV. (A = artery). Scale bar = 200 µm. **D** MIP of a z-stack from wholemount stained mesentery illustrates characteristic smooth muscle cell coverage (αSMA) and intraluminal valve formation (ITGα9, white arrowheads) in colMLV compared to LYVE-1 expressing capMLVs. Scale bar = 50 µm. **E** MIP of a confocal stack from a wholemount stained specimen at high magnification, showing capMLV exclusively composed of oak-leaf shaped capillary lymphatic endothelial cells (capLECs) characterized by discontinuous button junctions. Scale bar = 10 µm. **F** Confocal tile scan (MIP) showing the mutually exclusive expression of LYVE1 in capMLV and ACKR4 in colMLV. Scale bar = 50 µm. **G** UMAP presentation of cells sorted from the mesentery. Overall, 15 cell types were identified, including three LECs subtypes. **H** Violin plot representation of marker (PROX1, LYVE1, ACKR4) transcript levels in capMLECs and colMLECs isolated from the mouse mesentery. **I** Dot plot representing the single-cell transcriptome profiling, which defined the molecular characteristics of cell populations from adult mesentery. The color code indicates the scaled average expression level in each cluster while the dot size indicates the percent of cells in each cluster expressing the given gene.

Taken together, these results suggest the presence of a so far unappreciated, extensive lymphatic capillary network in the mouse mesentery that shares some characteristics with capLVs in other vascular beds, including blind ends, oak-leaf shaped LECs and intercellular button junctions. However, in stark contrast with other initial lymphatics, the capMLVs have an unusually elongated shape with only a moderate degree of branching and spread throughout the entire mesentery without connection with the coexisting network of colMLVs.

### Mesenteric lymphatic capillaries form perinatally in a *Vegfc*-dependent fashion from the valves of mature collecting lymphatic vessels

Having demonstrated the presence of the newly recognized capMLV network, we explored its origins, development and differentiation. Starting at E16.5, immature, LYVE1+ MLVs underwent maturation, adopting a collector phenotype. Maturation was characterized by a gradual loss of LYVE1 expression and onset of valve formation, indicated by the condensation of PROX1-high areas (Fig. S2 A arrowheads). Both processes were delayed in *Vegfc* heterozygous foetuses bearing a LacZ knock-in allele (*Vegfc*^+/LacZ^) (Fig. S2 B-D) (Karkkainen et al., 2004; Nurmi et al., 2015). *Vegfc*^+/LacZ^ fetuses expressed approximately 30 % less VEGF-C RNA in the mesentery than WT littermates, while RNA encoding the alternative VEGFR-3 ligand VEGF-D was not changed (Fig. S2 E-F). At E18.5, colMLVs in WT foetuses matured further, as evidenced by completion of LYVE1 downregulation and appearance of intraluminal valves (Fig. 2A, arrowheads). Both processes were delayed in *Vegfc*^+/LacZ^ E18.5 fetuses (Fig. 2 B). Surprisingly, at the same time, we noted the reappearance of LYVE1-positive vessels connected to the valve-forming territories of colMLVs in WT foetuses, marking the onset of capMLV development (Fig. 2C). Instead of the slim, LYVE1-positive capMLVs emerging from collector valves, we observed In *Vegfc*^+/LacZ^ foetuses wide-lumen MLV fragments and cysts, that have also been reported perinatally in *Vegfc*^lox/lox^; R26-CreER^T2^ pubs (Fig. 2 D, red and white arrowheads) (Nurmi et al., 2015).

**Figure 2.**
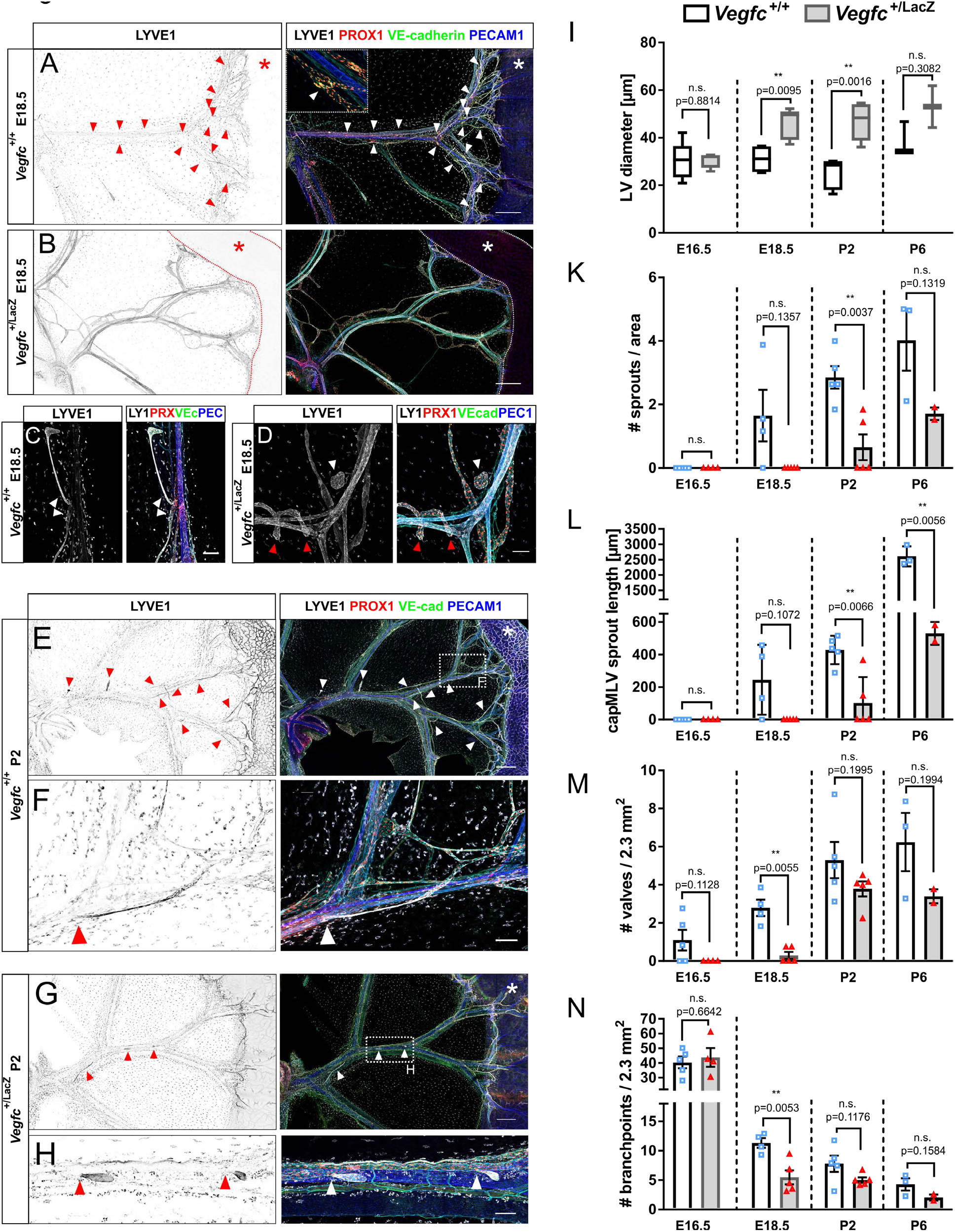
Mesenteric lymph vessel remodelling and maturation during peri-and early postnatal development are impaired in mice heterozygous for *Vegfc*. **A-D** Mesenteric wholemount immunostaining prepared from control (A, C; *n* = 4) or *Vegfc*^+/LacZ^ (B, D; *n* = 5) littermate foetuses (E18.5). Shown are MIPs of multi-tile z-stacks stained for LYVE1 (LY1), PROX1 (PRX), VE-cadherin (VEcad) and PECAM1 (PEC1). Arrowheads in (A) indicate intraluminal valve formation and hence maturation of colMLVs. The inset in (A) magnifies the valve area delimited by the white dotted box. White arrowheads in (C) mark newly forming LYVE1-positive capMLVs that appear in direct contact with valvular areas of mature colMLVs. Arrowheads in (D) denote blind ending LVs (red) and cyst-like structures formed by LECs (white) in *Vegfc*^+/LacZ^ mesenteries. Asterisks in (A, B) denote the position of the small intestine. Scale bars = 500 µm (A,B); 100 µm (C,D). **E-H** Mesenteric wholemount preparations stained for LYVE1, PROX1, VE-cadherin (VE-cad) and PECAM1 prepared from control (E, F; *n* = 5) or *Vegfc*^+/LacZ^ (G, H; *n* = 5) littermates at P2. Shown are multi-tile MIPs of z-stacks. The white boxed areas in E and G are magnified in F and H. White and red arrowheads highlight LYVE1-positive capMLV that emerge in direct contact with developing valvular territories in mature colMLVs. White asterisks in (E, G) identify the small intestine. Scale bars = 500 µm (E, G); 100 µm (F, H). **I-N** Quantification of lymphatic vessel characteristics in control and *Vegfc*^+/LacZ^ littermates. Stages of development are indicated. E16.5 control (n=5), E16.5 *Vegfc*^+/LacZ^ (n=4); E18.5 control (n=4), E18.5 *Vegfc*^+/LacZ^ (n=5); P2 control (n=5), P2 *Vegfc*^+/LacZ^ (n=5); P6 control (n=3), P6 *Vegfc*^+/LacZ^ (n=2). (K) Number of sprouts per lymph vessel area, (M) number of valves per viewfield of 2.3mm^2^, (N) number of branchpoints per viewfield of 2.3mm^2^. Data represent the mean ± s.d. Statistical significance is indicated above each group. Two-tailed unpaired Student’s *t*-test with Welch’s correction.

In WT pups, formation of capMLVs progressed between P2-P6, with the appearance of numerous LYVE1+ valvular extensions and their elongation (Fig. 2 E-F; Fig. S2 G). During this time window, we observed transient intraluminal valves inside newly formed capMLVs (Fig. S2 H, red arrowheads), but they had disappeared after the full segregation of capMLVs from colMLVs (Fig. 1).

MLV formation remained delayed also postnatally in *Vegfc*^+/LacZ^ pups (Fig. 2 G-H, Fig. S2 J-K) and *Vegfc* transcripts were approximately 30% reduced, while *Vegfd* transcripts were unaltered (Fig. S2 L-M). The retarded MLV development in the *Vegfc*^+/LacZ^ mice was associated with an increased LV diameter between E18.5 and P2 (Fig. 2 I) and, at all stages, in reduced formation of LYVE1+ sprouts (Fig. 2 K-L), valves (Fig. 2 M) and branches (Fig. 2 N).

The development of an independent, contiguous capMLV system closely aligned with mesenteric arteries was completed by P28, in the WT mice (Fig. 3A). In *Vegfc*^+/LacZ^ mesenteries, MLVs developed less well, with mild defects observed in colMLVs, including a slight dilation and, occasionally, imperfect valve maturation (Fig. 3 B-E). The capMLVs were severely hypomorphic, failed to elongate and remained attached to colMLVs as dilated stubs (Fig. 3 A – E). At this point in development, *Vegfc* transcripts in the mesentery were 50% reduced, while *Vegfd* transcripts were again not significantly altered (Fig. 3 F, G). The observed rudimentary capMLV-like vessels that retained valves (Fig. 3 K, L Fig. S3 A-E), consisted of oak-leaf shaped LYVE1+ LECs with discontinuous button-type junctions, which however were less defined than in WT capMLVs (Fig. 3 H-I).

**Figure 3.**
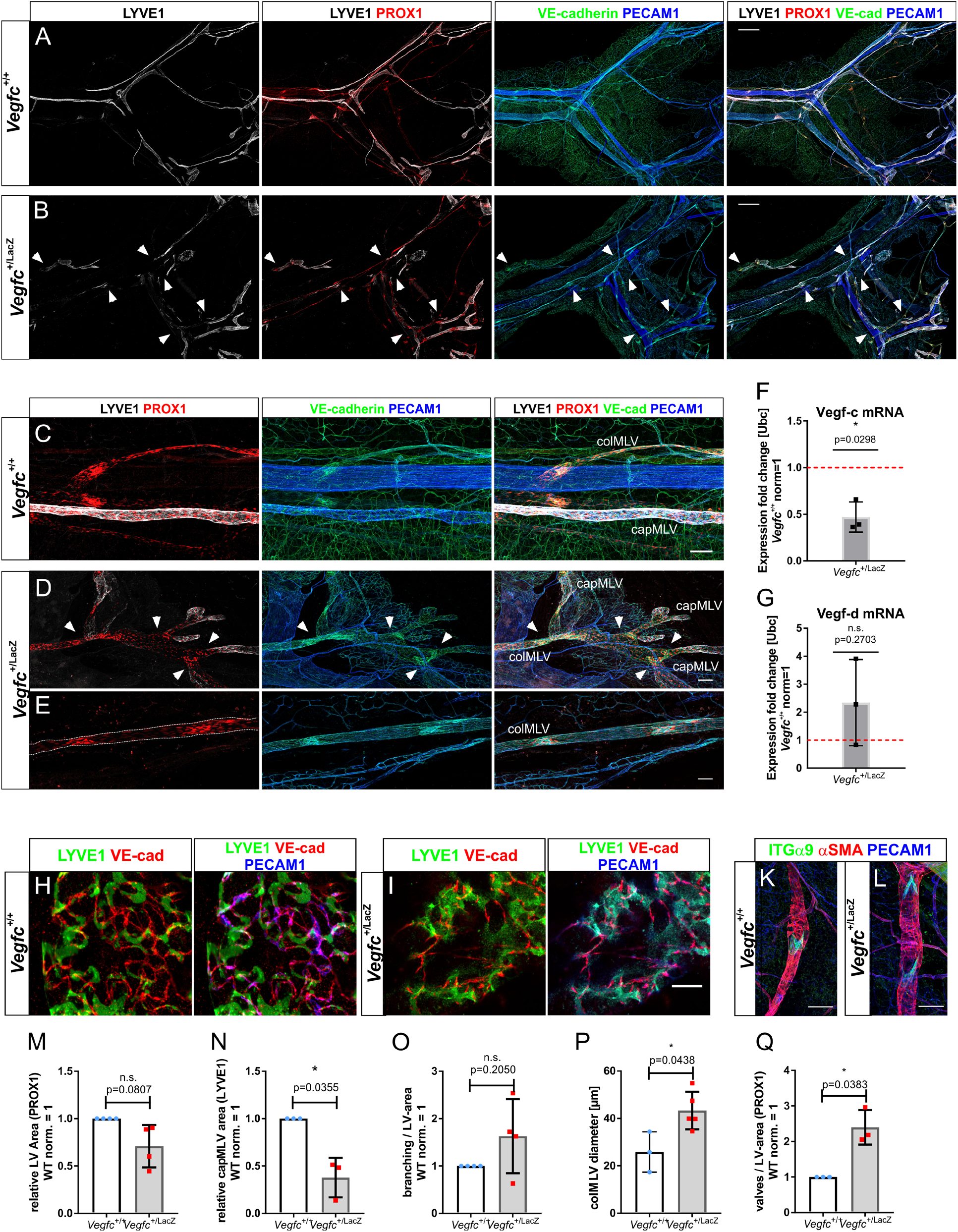
Mesenteric capillary network formation remains incomplete in adult mice heterozygous for *Vegfc.* **A-D** Representative mesenteric wholemount preparations from adult WT (A, C; *n* = 4) and *Vegfc*^+/LacZ^ (B, D-E; *n* = 5) littermates stained for the antigens indicated in color above each panel (VE-cad = VE-cadherin). A continuous capillary network, fully segregated from collectors, spanning the entire length between intestinal wall and lymph nodes is present in WT controls (A, C), while capillary LECs specify but fail to form into an independent, continuous network in *Vegfc*^+/LacZ^ mice (B, D-E). White arrowheads in (B, D) display sites of capillary formation that are incompletely segregated from mesenteric collectors. Shown are MIPs of multi-tile z-stacks. Scale bars = 500µm (A, B), 100µm (C-E). **F-G** Quantitative real-time (qRT)-PCR analysis of expression of the lymphangiogenic growth factors VEGF-C (F) and VEGF-D (G) in adult control and *Vegfc*^+/LacZ^ mesenteries. Individual samples were measured in triplicate. Relative expression (ΔΔ*C*t) of target genes was normalized to the control transcript *Ubc*. Vegfc^+/+^ expression levels were normalized to 1 (red dotted line). The data represent mean ± s.d (*n* = 3). Statistical significance is displayed above each graph. Two-tailed unpaired Student’s *t*-test with Welch’s correction. **H-L** Irregular oak-leaf shape and button junction formation of *Vegfc*^+/LacZ^ mesenteric capLECs (H, I), but normal smooth muscle cell coverage and valve formation in *Vegfc*^+/LacZ^ colMLV (K,L). Data are representative for four (control) and five (*Vegfc*^+/LacZ^) mesenteries. Shown are MIPs of single tile z-stacks. **M-Q** Quantification of the total lymph vessel area as defined by PROX1-expressing cells (M, *n*=4 control, *n*=4 *Vegfc*^+/LacZ^), capLV area (N, *n*=3 control, *n*=3 *Vegfc*^+/LacZ^), branch points per LV-area (O, *n*=4 control, *n*=4 *Vegfc*^+/LacZ^), colLV diameter (P, *n*=3 control, *n*=5 *Vegfc*^+/LacZ^) and number of valves per LV-area (Q, *n*=3 control, *n*=3 *Vegfc*^+/LacZ^). Values of WT normalized to 1 (M-O,Q). Data represent mean ± s.d. Statistical significance is indicated above each graph. Two-tailed unpaired Student’s *t*-test with Welch’s correction.

Thus, *Vegfc*^+/LacZ^ mice develop only incomplete, discontinuous capMLVs (Fig. 3M, N), that share properties with the pre-collector LVs of other vascular beds, including increased branching, increased diameter and retention of intraluminal valves (Fig. 3 O-Q).

### VEGF-C produced by arterial endothelial cells is indispensable for the formation of capMLVs

While we had established that the development of capMLVs is essentially depended on VEGF-C, their cellular source was unknown. Intrigued by the remarkable alignment of arteries and capMLVs, we employed the *Vegfc*^+/LacZ^ allele to localize VEGF-C producing cells and noted a strong LacZ/β-Gal staining reaction associated with mesenteric arteries (Fig. S4 A and B). In addition, transcriptomic and RT-qPCR analysis that BECs, fibroblasts, pericytes and smooth muscle cells were major sources of mesenteric VEGF-C (Fig 4 A-B; Fig. S4 C). To evaluate the importance of VEGF-C produced by arterial ECs vs other sources, we deleted *Vegfc* from *Vegfc^lox/lox^* in postnatal pups by using the tamoxifen-inducible BMX-CreER^T2^ allele (Fig.4 C), which selectively deletes in arterial ECs and collector LECs (Fig. S4 D, E). Because this *Vegfc*^lox^ allele is hypomorphic, *Vegfc*^lox/lox^ littermates and globally deleted *Vegfc*^lox/lox^; R26-CreER^T2^ pups were used as controls (Antila et al., 2017). Immunostaining revealed impaired capMLV development in hypomorphic *Vegfc*^lox/lox^ mesenteries, comparable to *Vegfc*^+/LacZ^ mice (Fig. 4D-D’, G). In the same specimen, colMLVs diameter, branching and valve formation was not altered (Fig. S4 F-F’). While only few LYVE1+ capMLVs were present in the the *Vegfc*^lox/lox^ mesenteries (Fig. 4 D-D’, G), tamoxifen-induced arterial deletion of *Vegfc* resulted in a total loss of capMLVs from the *Vegfc*^lox/lox^; BMX-CreER^T2^ mesenteries (Fig. 4 E, G). Meanwhile, collecting vessels had formed normally, and their branching and valve development remained unaffected (Fig. 4 H-L; Fig. S4 G-G’). Only postnatal global deletion in *Vegfc*^lox/lox^; R26-CreER^T2^ pups led to decay of the already established colMLVs (Fig. 4 F, G-L; Fig. S4 H-H’). After ubiquitous VEGF-C deletion, lymphatic valves were severely compromised. Although, clusters of PROX1 high cells were discernible, intact valve leaflets were not found (Fig. 4 F, K Fig. S4 H’). Despite these changes, further, clusters of LECs emerged from collecting vessels, leading to an increase in total LV area (Fig. 4 F; Fig. S4 H-L). In these LEC-clusters, LYVE1 was upregulated, suggesting loss of colLEC identity (Fig. 4 F). We noted also a gradual increase of the colMLV diameter with associated with VEGF-C levels (Fig. 4 H) and the development of chylous ascites in approx. 1/3^rd^ of the *Vegfc*^lox/lox^; BMX-CreER^T2^ mice and all *Vegfc*^lox/lox^; R26-CreER^T2^ mice, but not in the *Vegfc*^lox/lox^ mice (Fig. S4 I-L).

**Figure 4.**
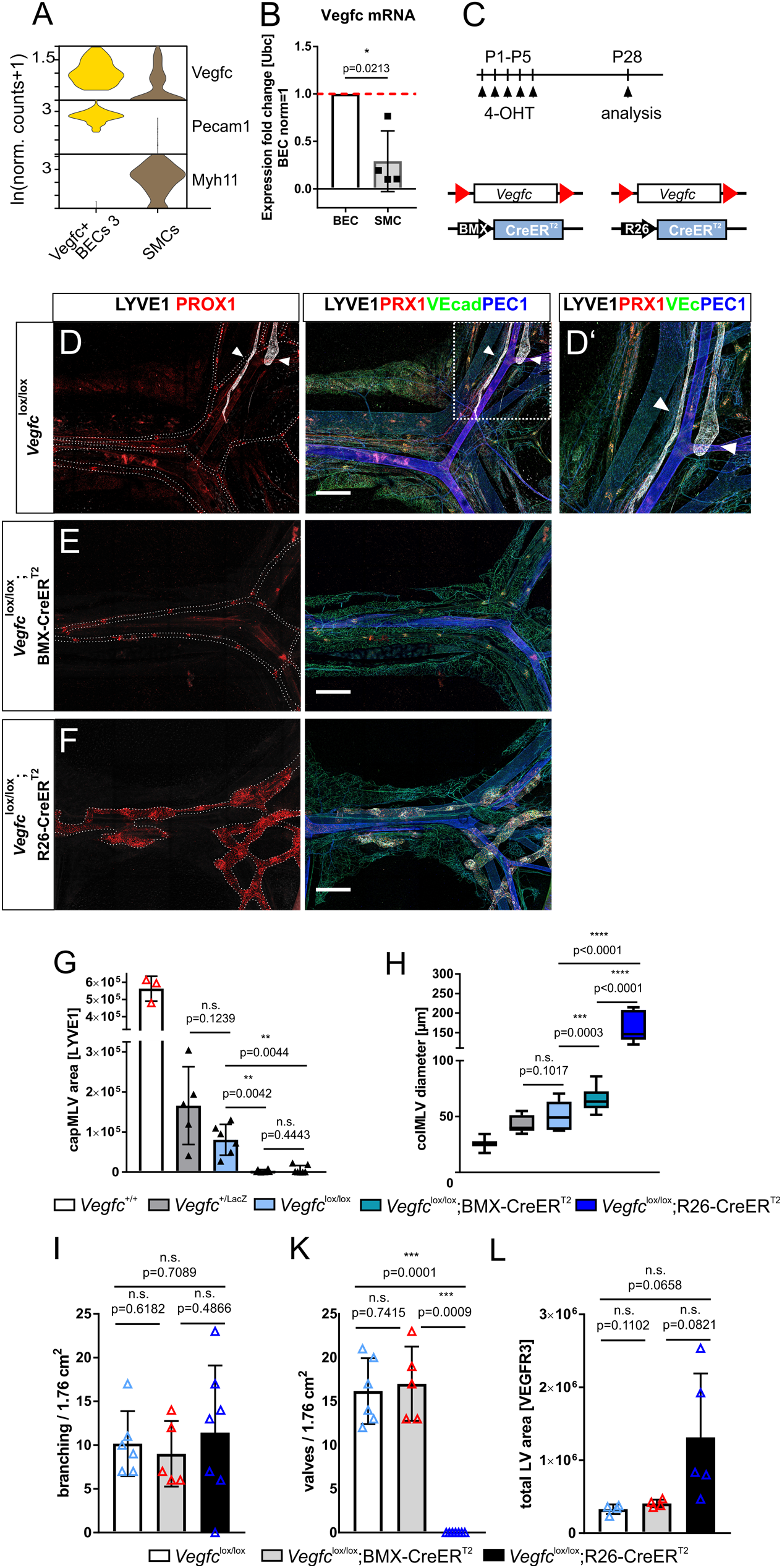
Deletion of *Vegfc* from arterial ECs-results in loss of mesenteric lymphatic capillary formation. **A** Violin plot representing Vegfc and PECAM1 / Myh11 transcript levels, the latter distinguishing BECs and SMCs in the mesentery. **B** Analysis of lymphangiogenic growth factor expression in BECs and SMCs (*n*=4) by qRT–PCR using specific TaqMan probes. For each experiment, samples from two mice of 14-16 wks were pooled and measured in triplicate. Relative expression (ΔΔ*C*t) of target genes was normalized to the control transcript *Ubc*. BEC expression levels were normalized to 1 (red dotted line). The data represent mean ± s.d. Statistical significance is indicated above the graph. Two-tailed unpaired Student’s *t*-test with Welch’s correction. **C** Schematic representation of the timing of 4-OHT administration and analysis and loci for postnatally induced arterial endothelial (BMX-CreER^T2^) or ubiquitous (R26-CreER^T2^) Cre-recombinase activation. **D - F** Representative multi-tile MIPs of mesenteric wholemount preparations from 4-OHT-treated *Vegfc*^lox/lox^ (*n*=6), *Vegfc*^lox/lox^; BMX-CreER^T2^ (*n*=6) and *Vegfc*^lox/lox^; R26-CreER^T2^ (*n*=7) mice at 28 days of age. Detected antigens indicated in color above the panels in E. (PRX1=PROX1, VEcad and VEc = VE-cadherin, PEC1 = PECAM1). White boxed area in D is enlarged in D’. Arrowheads in D, D’ denote capMLV in *Vegfc*^lox/lox^ mesenteries, while capMLVs were not detected in 4-OHT-treated *Vegfc*^lox/lox^; BMX-CreER^T2^ (L) and *Vegfc*^lox/lox^; R26-CreER^T2^ (M) mesenteries. Scale bars = 500µm. **G - L** Quantification of the capMLV area (G, occupied by LYVE1+ cells) and colMLV diameter (H) in control (*Vegfc*^+/+^ *n*=3), *Vegfc*^+/LacZ^ (*n*=5), and 4-OHT-treated *Vegfc*^lox/lox^ (*n*=6), *Vegfc*^lox/lox^; BMX-CreER^T2^ (*n*=6) and *Vegfc*^lox/lox^; R26-CreER^T2^ (*n*=7) mesenteries at P28. **I - L** Quantitative analysis of branch points (I), the incorporation of valves (K) and the total LV area (L, VEGFR3) in 4-OHT-treated *Vegfc*^lox/lox^ (*n*=4 (I); 6 (K,L), *Vegfc*^lox/lox^; BMX-CreER^T2^ (*n*=4 (I); 5 (K, L)) and *Vegfc*^lox/lox^; R26-CreER^T2^ (*n*=5 (I); 7 (K,L) mesenteries at P28. **G - L** The data represent mean ± s.d. Statistical significance is indicated above each graph. Two-tailed unpaired Student’s *t*-test with Welch’s correction.

These results demonstrated that capMLVs were exquisitely sensitive to a reduction of VEGF-C levels and that loss of arterial EC-derived VEGF-C sufficed to prevent capMLV development. In contrast, colMLVs were insensitive to reduced VEGF-C expression, and only decayed upon full loss of VEGF-C. In this scenario, compromised lymph flow due to degeneration of intestinal lacteals may have contributed to the colMLV deterioration. In summary, we demonstrate a unique dependence of the capMLV network on the availability of arterially produced VEGF-C.

### CapMLVs do not transport dietary components but drain only the peritoneal cavity

Having demonstrated the anatomical position, unique characteristics and differential growth factor requirement of the novel capMLV network, we probed its function. To assess the contribution of capMLVs to transport of dietary components, we administered the fluorescent lipid Bodipy-FL C16 via oral gavage and harvested the mesentery four hours later. Fluorescent lipids were detected in colMLVs after gavage, but did not appear in capMLVs (Fig. 5A, B). Conversely, i.p. administered, fluorescently labelled anti-LYVE1 antibodies readily stained capMLVs in less than 15 min, demonstrating direct access of peritoneal fluid to capMLVs (Fig. 5 C, D), but failed to decorate colMLVs. These results indicate drainage of dietary components and peritoneal solutes via the two distinct LV networks.

**Figure 5.**
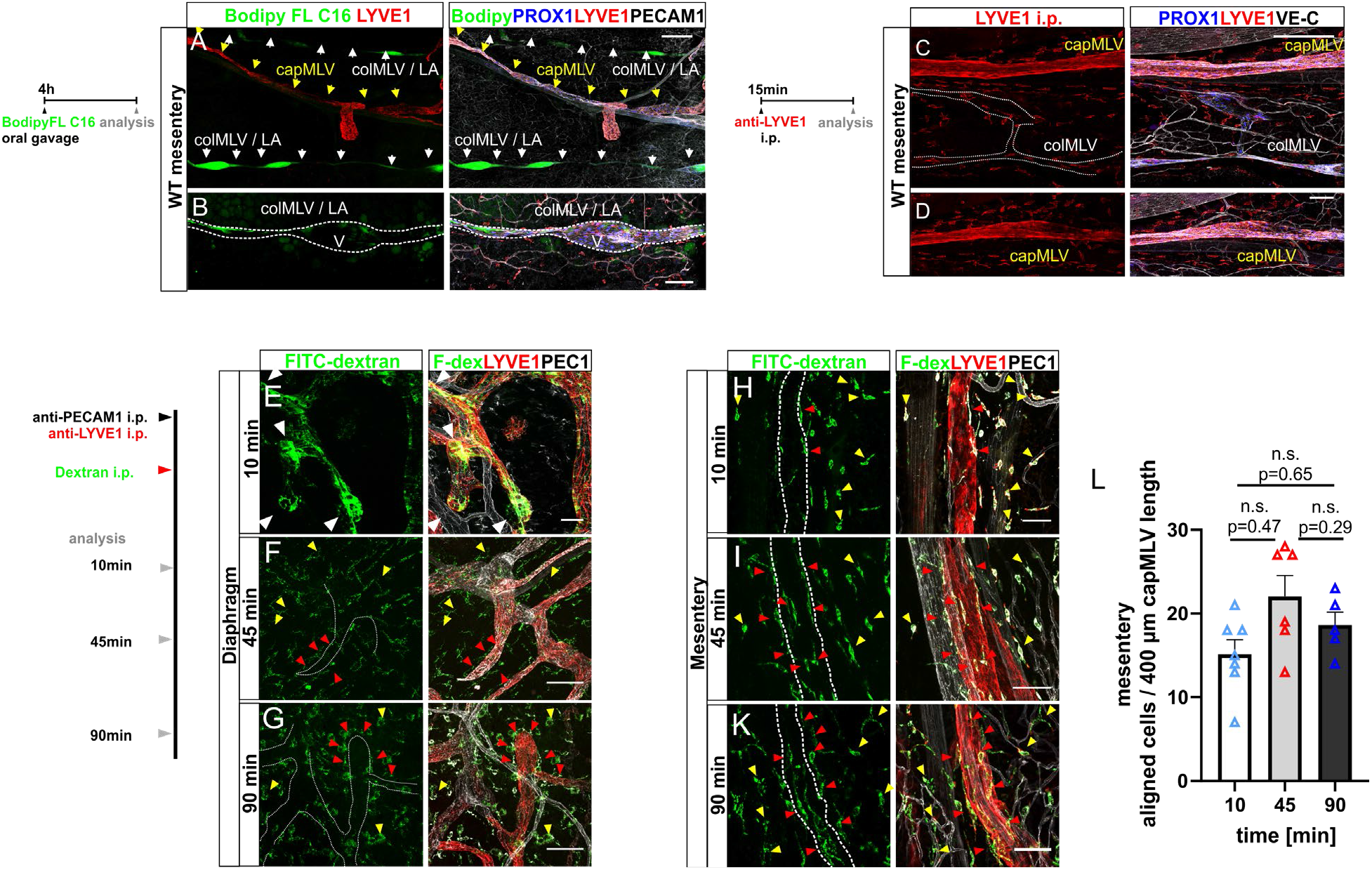
Peritoneal and intestinal transport functions are divided between mesenteric capillary and collecting lymphatic vessels. Schematics indicate time of administration and analysis for probing peritoneal lymph transport routes after oral gavage or i.p. injection. **A, B** Intestinal lymph transport route after oral gavage of 20 µg Bodipy FLC16 application. Multi-tile MIPs of wholemount immunostained mesentery four hrs after gavage (n=4). Colours of the fluorescent tracer and the detected antigens are indicated atop the panels. Yellow arrows denote a capMLV, white arrows colMLVs. V = valve, scale bars = = 100 µm (A), 50 µm (B). **C, D** Multi-tile MIPs of a wholemount immunostained mesentery 15 min after i.p. injection of LYVE1 antibodies (n=8). Colours of the detected antigens are indicated atop the panels, white outline = colMLV. Scale bars = 100 µm (C), 50 µm (D). **E - G, H - K** Peritoneal capMLVs constitute the preferential uptake and transport route for solutes (lysine fixable 70 kDa FITC-Dextran) after intraperitoneal injection, (E - G, diaphragm, n=6), (H - K, mesentery, n=6). Colours of tracer and detected antigens are indicated atop the panels. The data shows MIPs of tissues fixed *in situ* at the indicated times after i.p. tracer application. White arrowheads (E) highlight direct tracer uptake by blind-ended lymphatic capillaries. Yellow arrowheads point to dextran uptake by resident phagocytes and red arrowheads indicate the alignment and intravasion of phagocytes into capMLVs. Data represent MIPs of six individual experiments. Scale bars = 50 µm (E), 100 µm (F - K). **L** Quantitative analysis of phagocytes aligned over a length of 400µm to capMLVs at the indicated times following i.p. injection of FITC-dextran. Data represent mean ± s.d. Statistical significance is indicated above each graph. Two-tailed unpaired Student’s *t*-test with Welch’s correction.

Next, we injected tracers to identify the preferred routes and draining mechanisms for peritoneal solute transport. To visualize blood and lymphatic vessels *in vivo,* we injected a mixture of labelled anti-LYVE1 and anti-PECAM1 antibodies i.p., followed 20 min later by lysine-fixable 70 kDa FITC-Dextran injection i.p. to trace peritoneal fluid uptake. Mice were euthanized 10, 45 or 90 min after Dextran injection. To minimize intravascular tracer loss during dissection and preparation, we fixed the peritoneum *in situ* for 5 min with 0.4% PFA/PBS prior to dissection and wholemount preparation of diaphragm (Fig. 5 E-G) and mesentery (Fig. 5 H-K).

Among the anatomical sites analysed, the lymphatic lacunae of the diaphragm were preeminent in fluid transport. As early as 10 min after tracer application, FITC-dextran had entered capLVs in the center of the diaphragm (Fig. 5E, white arrowheads). Direct fluid uptake was no longer observed 45 min or 90 min after injection (Fig. 5 F-G). Instead, we found abundant tracer-positive, tissue resident LYVE1+ phagocytes that had the typical shape of recently described interstitial peritoneal macrophages (IPM) and localized near (Fig. 5 F-G, yellow arrowheads) or aligned closely to capLV (Fig. 5 F, G, red arrowheads) (Zhang et al., 2021). In contrast to the diaphragm, we did not observe direct uptake of FITC-dextran into *“mesenteric”* capMLVs (Fig. 5 H-K).

Already 10 min after injection, LYVE1+ IPMs in the mesentery had phagocytosed FITC-dextran (Fig 5 H-K, red & yellow arrowheads). Subsequently, they closely aligned with and entered capMLVs (Fig. 5 H-K red arrowheads, Fig. S5 A-B white arrowheads). The association of LYVE1+ IPM and capMLVs peaked at 45 min after tracer application (Fig. 5 L). LYVE1+ IPM of the mesentery were dendriform and localized between the mesothelial sheets of the mesenteric vascular and avascular areas. They were clearly distinct from peritoneal fluid macrophages (PFM), which were more round and localized towards the cavity surface atop of the mesothelium (Fig. S5 C). As capMLVs strongly express CCL21 (Fig. 1 I), and we detected CCR7+ cells in the proximity of capMLVs (Fig. S5 D, white arrowheads), we speculated that guidance to capLVs may rely on the CCR7 - CCL21 signalling axis. Reanalysis the LYVE1pos and LYVE1neg macrophages from two published data sets containing scRNAseq data from murine mesenteric cells (GSE102665) and omental macrophages (E-MTAB-8593) revealed that LYVE1+ IPM do not express CCR7 (Fig. S5 E, F) (Zhang et al., 2021). Furthermore, since i.p. injections engage capLVs of the visceral and parietal serosa, we detected tracer in mediastinal, coeliac and inguinal LNs, but never in mesenteric LNs (Fig. S5 G-K), reinforcing the notion of two distinct draining routes for orally ingested and peritoneal antigens.

### CapMLVs drain to mediastinal lymph nodes

The finding of separate transport routes for intestinal and peritoneal matter, engaging colMLVs and capMLVs respectively, prompted us to identify the capMLV-draining LNs. However, tracer uptake by capLVs in both visceral and parietal serosa did not allow identification of the specific LNs draining the capMLVs. Several studies have identified MLNs as the colMLVs-draining LNs (Czepielewski et al., 2021; Friess et al., 2022; Gautreaux et al., 1994; Houston et al., 2016; Worbs et al., 2006). The MLN chain, located at the branch point of superior mesenteric artery and abdominal aorta, is rich in fat, which impeded a microsurgical analysis of vessel routing. To analyse the track of capMLVs in this dense tissue convolute, we resected the intact organ complex consisting of duodenum, jejunum and ileum with the mesentery including its vasculature, central LNs and fat body attached (Fig. S6 A-D). Following wholemount immunostaining and tissue clearing of the ensemble, we used light sheet microscopy to generate image stacks, from which 3D tissue renderings with minimally disturbed vessel architecture were digitally reconstructed (Fig. 6 A-E, Fig. S6 A-C’’). We distinguished ACKR4 positive colMLVs from ACKR4 negative capMLVs by taking advantage of a knock in allele, which expresses eGFP under control of the ACKR4 promotor (*Ackr4^GFP^*) (Friess et al., 2022; Heinzel et al., 2007). Strong VEGFR-3 immunoreactivity of the regularly spaced intraluminal valves further confirmed colMLV identity (Fig. 6 B’). In volume reconstructions of the wholemount preparations, GFP+, valve-containing colMLVs were readily distinguishable from LYVE1+, capMLVs (Fig 6 A-E’). ACKR4+ LECs, lining the ceiling of the MLN subcapsular sinus, connected directly with colMLVs (Fig. 6 B-B’, D – E, white arrowheads). We never observed such connections between MLN subcapsular sinus and LYVE1+ capMLVs, which instead bypassed the MLN chain and tightly followed the superior mesenteric artery (Fig. 6 A-E).

**Figure 6.**
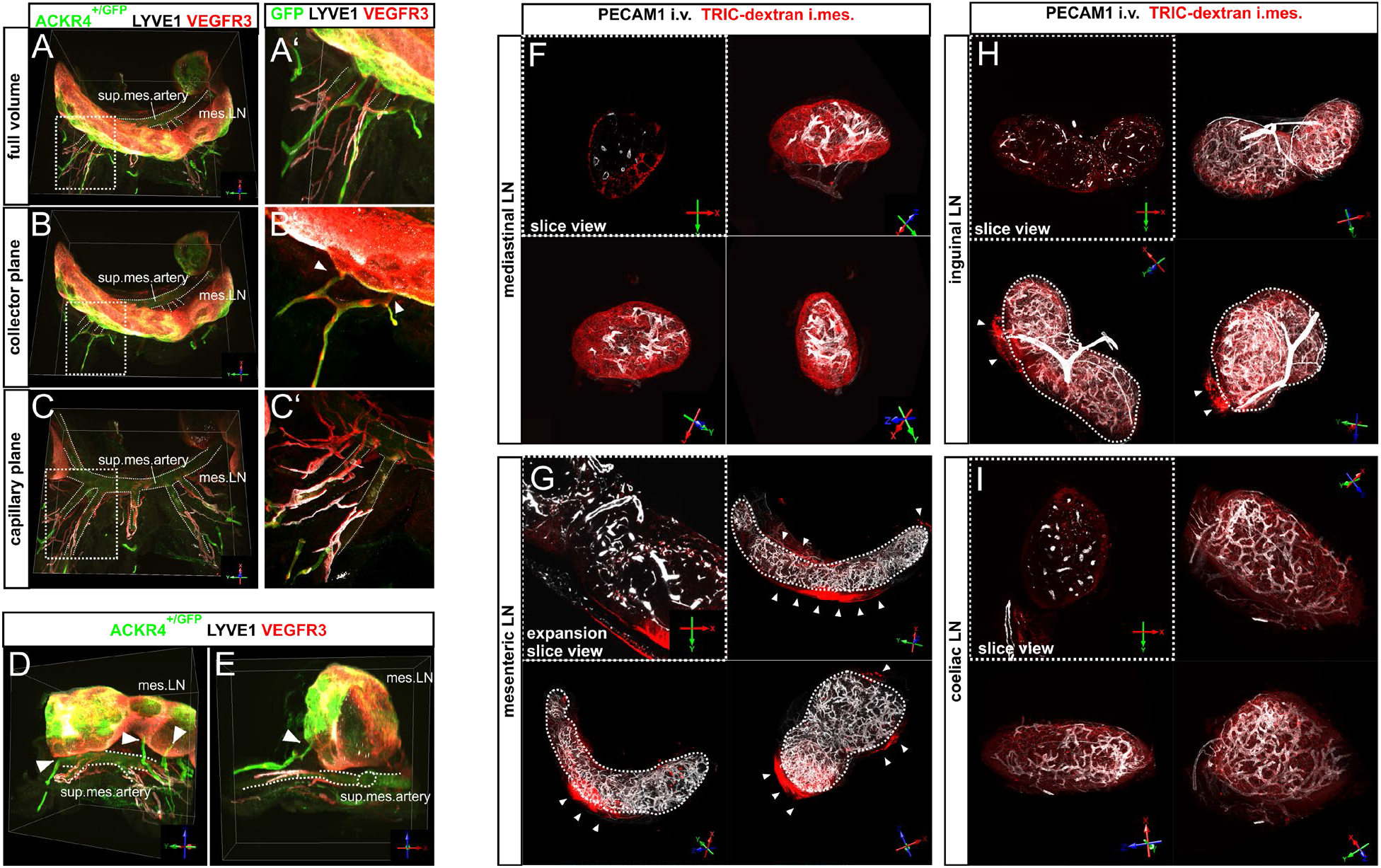
Identification of the draining lymph nodes of mesenteric lymphatic capillaries. **A - C** Projections of digital volume reconstructions of full wholemount preparations (shown in Fig.S6, D) after immunostaining for the indicated antigens followed by optical clearing for light sheet microscopy. Left row: reconstruction of the full volume (A) or two selective axial slices (B + C), volumes delineated by the white boxes are shown higher magnified in the right row (A’ – C’). **D + E** Volume renderings demonstrating direct drainage of GFP+ (ACKR4 expressing) mesenteric colMLVs into the mesenteric lymph nodes (MLNs), situated atop the superior mesenteric artery. Note the position of the LYVE1+, VEGFR3+ capillaries following the artery beneath the node. Lymph node LECs at the roof of the capsular sinus are also ACKR4+ (GFP+). **A – E** White arrowheads (B’, D, E) point to ACKR4+ colMLVs that drain to the MLNs. The white dotted line outlines the superior mesenteric artery and its branching arteries supplying the small intestine. The data show representative images of four individual experiments. **F - I** Application of the tracer TRITC-dextran via intramesenteric injection to identify the draining lymph nodes of capMLVs (n=7). The blood vasculature had been pre-labelled by i.v. application of PECAM1 antibodies. 20 min after TRITC-dextran application, LNs were prepared and cleared for light sheet microscopy. Shown are digital sections of 2 µm thickness (white dotted rectangles) and full volume renderings (three axial directions) of a representative mediastinal, mesenteric, inguinal and coeliac node. White dotted lines in H, I outline the mesenteric and inguinal LN. White arrowheads highlight tracer accumulated on the outside of the LN capsule. sup.mes.artery - superior mesenteric artery; mes.LN – mesenteric lymph nodes; i.mes –intramesenteric injection, i.v. – intravenous injection.

Having excluded MLNs as draining LNs for capMLVs, we focussed on the gastric and pancreatoduodenal LNs, located alongside the coeliac trunk (Fig. S6 D), but the 3D tissue rendering analysis again revealed no direct nexus to capMLVs. Instead, capMLVs followed the abdominal aorta and exited the peritoneal cavity (Fig. S6 B-C’’ white arrowhead). Attempting to further trace capMLVs beyond the diaphragm, we reached the limits of the wholemount preparation approach. Since preferential drainage via the capLVs of parietal serosa and diaphragm rendered i.p. tracer injections uninformative, we resided to microinject fluorescent tracer (TRITC-dextran) *“intramesenterically”* into the externalized mesentery and repositioned the intestinal loop until further analysis. After 20, 45 or 90 min mesenteric, mediastinal, coeliac and inguinal LNs were dissected, fixed and either analysed as cryosections (Fig. S6 E-G) or optically cleared for light sheet microscopy (Fig. 6 F-I). In agreement with our previous analysis, tracer did not accumulate in the MLNs. A rim of tracer lining the outside of the MLN capsule (Fig. 6 G, arrowheads) was likely due to flooding along the mesenteric vascular bundles reaching the outer perimeter of the MLNs. In stark contrast, already 20 min after its application, TRITC-dextran was found in the cortex, sub-capsular and medullary sinues of mediastinal LNs (Fig. 6 F, Fig. S6 E-G), indicating rapid uptake and transport via capMLVs. Compared to the uptake to mediastinal LNs, distinctly less tracer reached the coeliac and inguinal LNs (Fig. 6 H, I).

In summary, injection experiments probing capMLV function confirmed our previous observation of strictly separate lymph vessel routes for dietary and peritoneal antigens. Surprisingly, the latter were preferentially transported from capMLVs to the mediastinal LNs and thereby reached a distinct site of for presentation to the adaptive immune system.

### Cells in the avascular planes of the mesentery juxtapose capMLVs and express ACKR4, implying the CCL21-CCR7 chemokine - receptor axis in antigen transport to capMLVs

Having established preferential drainage of capMLVs into the mediastinal LNs, we investigated possible mechanisms supporting antigen trafficking to the capMLVs in the vascular zone of the mesentery (Fig. S7 A). ScRNA-Seq analysis of capMLEC and colMLEC populations identified CCL21 as the only chemokine prominently expressed on the ACKR4-negative capMLECs (Fig. 7 A, Fig. S7 A-B). We confirmed CCL21 expression on capMLVs by immunostaining (Fig. 7B, Fig. S5 C), while ACKR4+ colMLECs lacked chemokine expression (Fig. 7A-B, Fig. S7 B). The CCL21 - CCR7 chemokine – chemokine receptor pair directs leukocyte traffic to LNs (Arasa et al., 2021; Bryce et al., 2016; Forster et al., 2008; Heinzel et al., 2007; Russo et al., 2016). CCL21 availability is controlled by the atypical chemokine receptor 4 (ACKR4), that scavenges CCL21 and cognate chemokines from the extravascular space, of LN subcapsular and splenic peri-marginal sinuses and from colMLVs (Friess et al., 2022; Heinzel et al., 2007; Ivanov et al., 2016; Ulvmar et al., 2014). After tracer injection, we noted that the intermesothelial space of the avascular zone contained abundant, tracer-loaded, mainly LYVE1+, CCR7+ IPMs, interspersed with non-phagocytotic ACKR4 expressing cells (Fig 7 E, Fig. S7 C-D), that also expressed PDFGRα and vimentin, but not LYVE1 and CCR7 (Fig.S7 C-D). ACKR4+ cells were massively depleted or completely absent within a rim of approx. 100 to 400 µm at the border between the vascular bundles and the avascular zone (Fig. 7 C-D, F). The formation of a chemoattractant CCL21 gradient is reinforced by ACKR4 via CCL21 scavenging from the extravascular space, as shown in multiple studies. We interrogated the functional requirement for ACKR4, using homozygous ACKR4^GFP/GFP^ knock-in mice, which are ACKR4-deficient but express eGFP instead. Unexpectedly, we found that upon ACKR4 deletion, eGFP+ cells were no longer excluded from the border between avascular and vascular zone, but had invaded the vascular zone, were ACKR4^+/GFP^ cells had not been detected before (Fig. 7 F-G). This indicated that the positioning of the intermesothelial cells might be dependent on ACKR4 expression and potentially, chemokine signalling. Furthermore, trafficking of LYVE1+, CCR7+ phagocytes to the capMLVs was strongly impaired after ACKR4 deletion (Fig. 7 H-I), suggesting an essential role of ACKR4 in the movement of phagocytes to capMLVs.

**Figure 7.**
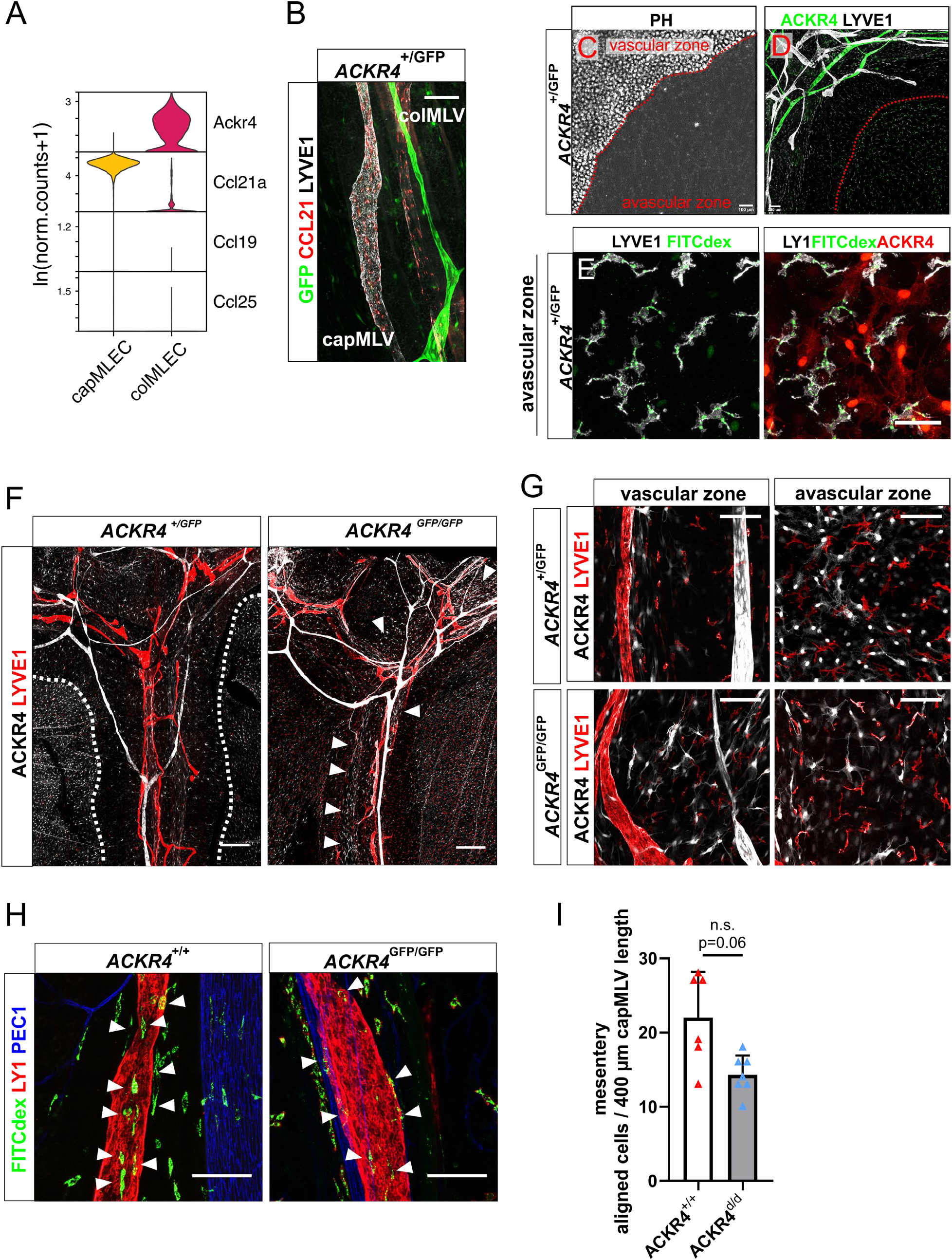
The atypical chemokine ACKR4 contributes to the recruitment of active phagocytes to capMLVs **A** Violin plot of the transcript levels of selected markers in mesenteric capMLECs and colMLECs. **B** Multi-tile MIP of an immunostained mesentery wholemount preparation discriminating capMLVs and colMLVs based on CCL21 and LYVE1 expression but absence of ACKR4, which is restricted to colLVs. Scale bar = 100µm. **C – D** Multi-tile MIPs of an immunostained ACKR4^+/GFP^ mesentery wholemount preparation, antigens indicated in colour above each panel. The red dotted line in **C** indicates the transition between the vascular and avascular zones, while the area delimited by the red dotted line in **D** contains ACKR4+ resident cells limited to the avascular zone. **C** PH = phase contrast identifies the adipose tissue covering the mesenteric vessel bundle. Scale bar = 100µm. **E** MIP of the avascular zone (ACKR4^+/GFP^ mesentery) 15 min after intraperitoneal FITC-dextran injection. Immunostained antigens are depicted in colour above each panel. Scale bar = 50µm. **F, G** Multi-tile MIPs of a mesenteric vessel branch in an ACKR4^+/GFP^ or ACKR4^GFP/GFP^ (ACKR4-deficient) mouse stained for capMLVs and colMLVs. **G** highlights the absence / presence of ACKR4+ cells within the vascular and avascular zones of the mesentery. Scale bar = 500 µm (F), 100 µm (H). **H** MIPs of ACKR4^+/+^ and ACKR4^GFP/GFP^ mesenteries fixed 45 min after i.p. application of the tracer FITC dextran. White arrowheads indicate FITC dextran uptake by resident phagocytes, which aligned with or entered capMLVs. The data are representative of six individual experiments. Scale bars = 100 µm. **I** Quantitative analysis of phagocytes aligned to 400µm length of capMLVs in the mesentery of ACKR4^+/+^ and ACKR4^GFP/GFP^ mice, 45min following FITC-dextran i.p. injection. The data represent mean ± s.d. Statistical significance is displayed above the graph. Two-tailed unpaired Student’s *t*-test with Welch’s correction.

Taken together, we show that the atypical chemokine receptor ACKR4, which is expressed on interstitial cells in the avascular zone of the mesentery, is involved in the trafficking of LYVE1+ IPM to colMLVs. Unexpectedly, we also found that ACKR4 affects the positioning of intermesothelial resident cells.

## Discussion

Here we reveal, for the first time a network of LVs in the mouse mesentery that is anatomically and functionally distinct from the well-established system of mesenteric LVs draining the gut into the MLNs. The existence of two independent, mesenteric LV beds with dissimilar properties sharply contrasts with other anatomical sites, all containing a sole network with a hierarchical LV architecture (Petrova and Koh, 2018; Sabine et al., 2016; Schulte-Merker et al., 2011). Such unique lymphatic status of the mesentery has important functional implications. Two separate drainage routes provided by the capMLVs and colMLVs into the mediastinal LNs and MLNs, respectively, allow for separation of peritoneal immunity and oral tolerance alternatively required to confront alimentary and pathogenic antigens.

During development of the mesentery, LECs differentiate from veins at the mesenteric root and simultaneously, from hemogenic endothelium to jointly form the primitive, LYVE1+ LV plexus (Sabin, 1902; Stanczuk et al., 2015; van der Putte, 1975). Before birth, this vascular network starts to remodel into mature colMLVs, which is associated with LYVE1 downregulation and the onset of valve maturation. We found the origin of capMLV formation in new LYVE1+ vessels evolving perinatally from the valve areas of colMLVs. The capMLVs elongated and remodelled during the first four postnatal weeks with valves formed transiently until complete segregation of the two LV systems. Similar postnatal development of LVs was described in the liver and meninges (Antila et al., 2017; Bobe et al., 2022). As in many other LV beds, VEGF-C was the major indispensable developmental lymphangiogenic growth factor also in mesenteric LV development (Haiko et al., 2008; Karkkainen et al., 2004; Nurmi et al., 2015). In *Vegfc* heterozygous mice, embryonic capMLV specification was delayed and incomplete zipper to button transformation impacted the development of well-defined oak-leaf shaped LECs. The persistent hypoplastic capMLVs of reduced length failed to fully remodel into an independent capMLV network. This resembled the description of the hypoplastic dermal lymph vasculature, intestinal lacteals and colMLVs in postnatally *Vegfc* deleted mice (Nurmi et al., 2015), although the colMLVs ultimately formed, indicating differential VEGF-C requirements in the different MLV types. The predominant transcription of VEGF-C in arterial ECs and SMCs and the close alignment of capMLVs and arteries suggested that arterial-derived VEGF-C is required for capMLV growth, positioning and maintenance. This was confirmed by VEGF-C deletion in arterial BECs, which eliminated capMLVs but not colMLVs. Already long time ago, a study highlighted the proximity of initial lymphatics to feeding arteries in the connective tissue located between the mesenteric vessel bundles in cats (Schmid-Schonbein, 1990). The BMX-Cre also induced gene deletion in colLVs, as expected, but this had no consequences because they lacked VEGF-C expression. The strong dependency on VEGF-C levels resembles findings in postnatally forming meningeal LVs and suggests a special requirement for VEGF-C in postnatally developing LVs (Antila et al., 2017).

The well-established colMLVs transport fat-soluble dietary components, and soluble or cell-borne antigens from the intestinal lacteals and the submucosal lymphatic network in the gut to the MLNs (Choe et al., 2015; Czepielewski et al., 2021; Unthank Joseph L, 1988). This process is fundamentally required for the induction and maintenance of oral tolerance (Huang et al., 2000; Worbs et al., 2006). The alternative, route described here, transports peritoneal fluid, antigens and antigen-laden cells into mediastinal LNs, establishing these as hitherto not recognised important elements of peritoneal immunity. In line with the here described capMLVs network, the lymph vessel beds of the meninges (Ahn et al., 2019; Antila et al., 2017) heart (Brakenhielm and Alitalo, 2019) and liver (Bobe et al., 2022) have been recently reported to be formed nearly exclusively of capLVs. This reveals that capLVs alone can act as an independent LV network within organs, where uptake of tissue fluid and immune cells is required, while drainage into afferent colLVs likely takes place outside of the organ. Presence of mesenteric capMLVs, based on PROX1 and LYVE1-immunoreactivity and oak-leaf shaped LECs with discontinuous junctions has been reported in humans, cats and rats, but their precise architecture, completely autonomous nature, and unique draining properties by-passing the MLNs have not been described previously (Murfee et al., 2007; Roncati et al., 2018; Sweat et al., 2012).

Unlike in the LVs of the diaphragm, we observed that tracer was drained via the capMLVs only within phagocytotic LYVE1+ cells. Although we cannot exclude that we missed direct, rapid fluid transport through capMLVs, abundant presence of stomata in the mesothelium atop the lymphatic lacunae of the diaphragm and the periodic alternation of positive and negative pressure due to breathing provide an explanation for the higher efficacy of lymphatic fluid transport in them. In the mesentery, numerous stomata are located in the mesothelium covering the avascular zones. These stomata allow entry of peritoneal fluid to the submesothelium, they provide a significant surface for antigen uptake and sampling by phagocytes (Abu-Hijleh et al., 1995; Bellingan et al., 2002; Wang et al., 2010), but are unlikely to exchange fluid with the vessel bundles of the vascular zones by interstitial convection. We found rapid phagocytosis of i.p. injected soluble tracer by LYVE1+ interstitial macrophages, supporting the concept of direct fluid access to the interstitial space of vascular and avascular zones.

So far, four distinct peritoneal macrophage populations have been described. These include two present in suspension in the abdominal cavity, large peritoneal macrophages (LPMs) and IRF4-dependent small macrophages. Two additional populations of interstitial peritoneal macrophages (IPMs) have been described recently, characterized by LYVE1^hi^ MCHII^lo-hi^ CX_3_CR1gfp^lo/-^ and the LYVE1^lo/-^ MCHII^hi^ CX_3_CR1gfp^hi^markers (Zhang et al., 2021). Both populations evenly distributed across the surface of the vascular and avascular zones of the mesentery. As judged by their LYVE1 expression, shape and spatial distribution, the interstitial LYVE1+ macrophages we describe here are IPMs, which are predominantly derived from embryonic precursors, long lived and distinct from the free floating peritoneal macrophages. Potential distinct functions of the sub-populations of peritoneal macrophage in phagocytosis and migration via LVs to the draining lymph nodes remain to be defined. Following i.p. injection of tracer, LYVE1+ IPM in the vascular and bordering avascular zones aligned with and entered capMLVs. This process was supported by ACKR4+ mesenchymal cells, localized within the interstitium of the avascular zone.

High expression of the hyaluronan receptor LYVE1 and the chemokine CCL21 are characteristic for capMLVs in distinction from colMLVs (Mäkinen, 2005). Both molecules have been implicated in guidance and entrance of CCR7+ DCs and CD4+ effector memory T cells from the periphery to LNs (Arasa et al., 2021; Bryce et al., 2016; Forster et al., 2008; Heinzel et al., 2007; Russo et al., 2016; Ulvmar et al., 2014). Local removal of CCL21 by ACKR4, expressed by multiple stromal cells, is a critical component of CCL21-mediated immune cell guidance that operates in ECs of the peri-marginal sinus of the spleen, in keratinocytes and in LECs lining the ceiling of the LN subcapsular sinus and those in afferent lymphatic collectors (Friess et al., 2022; Takeda et al., 2019; Ulvmar et al., 2014; Werth et al., 2021). In these microenvironments, ACKR4 has been shown to scavenge CCL19 and/or CCL21, reducing their availability and contributing to the shaping and stabilizing functional chemokine gradients and patterns (Bryce et al., 2016; Ulvmar et al., 2014; Werth et al., 2021). Under steady state conditions, macrophages do not express CCR7, which also holds for LYVE1+ IPMs, however, it can be induced under pathological conditions (Kwiecien et al., 2019; Tateyama et al., 2009). Recent studies have suggested that CCR7+ DCs can guide CCR7-, CCR5+ macrophages by CCL5 production to the draining LNs (Rawat et al., 2023).

The molecular profile of capLVs, their button junctions and expression of the chemoattractant CCL21 support peritoneal fluid and tissue-resident intermesothelial immune cell transport by these vessels. Our finding of a singular capillary lymphatic network that works in parallel with the collecting lymphatics in the mesentery provides an anatomical basis for a unique peritoneal immune surveillance distinguishing food antigens and peritoneal pathogens. This should have far-reaching implications for disease pathogenesis, when thinking for example the fact that homeostatic immune cell trafficking is prone to hijacking by cancer cells.

## Acknowledgments

We thank Stefan Volkery and Malte Stasch for excellent help with microscopy, Barbara Waschk and Jörn Prinz for help with genotyping. We thank Dietmar Vestweber for sharing antibodies and for fruitful discussions, Serge van de Pavert and Noelia Alonso Gonzalez for critically reading the manuscript. We gratefully acknowledge funding by the Deutsche Forschungsgemeinschaft (DFG, German Research Foundation) – CRC-1348/1 – 386797833 to FK and ER and SFB1450/1–431460824 to NK and FK.

## Author Contributions

ER and FK conceived the project, designed experiments, analysed data and wrote the manuscript; ER, NK, SF, AGL, MS, TW designed and performed experiments, analysed and interpreted data; RA provided reagents and experimental support; KA, TVP, AR provided mice and reagents, enabled experimentation and contributed to the manuscript.

## STAR Methods

### Resource availability

#### Lead Contact

Further information and requests for resources and reagents should be directed to and will be fulfilled by the Lead Contact, Friedemann Kiefer.

#### Materials Availability

This study did not generate new unique reagents.

#### Data and Code Availability

The single cell RNA sequencing data analyzed in this study was published by (Gonzalez-Loyola et al., 2021) and is available in NCBI’s Gene Expression Omnibus (GEO) under accession number GSE155954. RNA sequencing data of single cells including macrophages was also publicly available in GEO, accession number GSE102665, and in ArrayExpress, accession number E-MTAB-8593. All R packages and code used were published by their respective authors; no new code was developed specifically for this study.

## Experimental model and subject details

### Animal experiments

Mice were kept under conventional conditions in IVC-cages and ventilated racks at 22 °C and 55 % humidity with a light-dark cycle of 14:10 h. Regularly mice were tested for infectious disease including bacterial or fungal infections and parasites. Mice of indicated strains were analyzed at the age of postnatal day 2 (P2) to 28 weeks. Foetuses were analyzed at developmental stages between E16.5 and E18.5. Embryonic staging was determined by the day of the vaginal plug (E 0.5). Cre-negative littermate mice or embryos served as controls. Other control mice were matched by genetic background, sex, and age. All Procedures involving animals were carried out in strict accordance with the local animal protection legislation and were approved by the corresponding ethics committees. Approval is either documented in protocol TVA 81-02.04.2022.A226 (Münster) or the lab animal licence ESAVI/6036/04.10.07/2016 (Helsinki).

## Method Details

### Induction of Cre-mediated deletion of conditional KO mice

For *in vivo* induction of Cre recombinase activity, *Vegfc*^lox/lox^, *Vegfc*^lox/lox^; BMX-CreER^T2^ and *Vegfc*^lox/lox^; R26-CreER^T2^ mice were injected intragastrically at P1-P5 with 2 µL of 25 mg/mL Z-4-hydroxytamoxifen (Z-4-OHT) dissolved in 97 % ethanol. Msn and intestinal samples were analyzed at P28.

For the analysis of BMX expression R26-tdRFP; BMX-CreER^T2^ mice were injected intraperitoneally with a single suboptimal dose of Z-4-OHT (0.3 mg at P1) dissoved in peanut oil and analyzed at P28.

### Oral lymphangiography

Mice at P28 to 10 weeks of age were fed 20 µg 4,4-Difluoro-5,7-Dimethyl-4-Bora-3a,4a-Diaza-s-Indacene-3-Hexadecanoic Acid (Bodipy-FL C_16_) diluted in peanut oil by oral gavage. Four hours later, mesenteries were harvested and subjected to wholemount immunostaining.

### Intraperitoneal injection of antibodies

To analyze uptake of intraperitoneal fluid by msn capillaries, mice at P28 were injected intraperitoneally with 0.75 mg/kg LYVE1 antibody (R&D Systems, clone 223322) coupled to Alexa 594 diluted in 200 µL PBS. Mesenteries were harvested 15 minutes later and wholemount immunostained for vessel type-specific antigens.

### Identification of peritoneal lymphatic routes and draining lymphnodes

To analyze lymphatic vessel uptake and draining LN of intraperitoneally injected fluid, mice at P28 were injected with 200 µg lysine-fixable FITC-dextran (70 kDa), diluted in 200 µL PBS. Co-Injection of 0.75 mg/kg LYVE1 antibody (R&D Systems, clone 223322) coupled to Alexa 594 and 20 µg PECAM1 antibody (clone 1G5 + 5D2, Wegman et al., 2006) coupled to Alexa 647 identifies lymphatic and blood vessels. Tissues were pre-fixed with 0.4 % PFA/PBS *in situ* prior to preparation a full fixation with 4 % PFA/PBS. Mesenteries were harvested after 10, 45 or 90 minutes and directly mounted for microscopy.

### Intramesenteric injections

To specifically trace capMLV drainage into LN, we developed a surgery to expose the mesentery and to inject tracers directly into the mesentery. Mice of 8-16 weeks received an i.p. injection of 0.04 mg Fentanyl / 4 mg Midazolam / 1 kg bodyweight and were anaesthetised with 4 % v/v Isoflurane/O2 (Abbott Animal Health) and maintained with 2% v/v Isoflurane/O2. To label blood vessels, 15 µg PECAM1 (clone 1G5+5D2, Wegman et al., 2006) antibody coupled to Alexa 647 in 50 µL total volume (0.9 % NaCl) were injected retrobulbar. Animals were fixed on a heatpad, the abdominal cavity was opened by vertical incision of approx. 1cm and individual intestinal loops with mesenteric branches were exposed. TRITC-dextran/0.9% NaCl in 50 µL injection volume were injected into 4-6 individual mesenteric branches close to the intestinal attachment. The intestine was repositioned and the abdominal cavity was sewed. Lymphatic drainage was analyzed after 20, 45 and 90 min. Therefore, lymphnodes (inguinal, mesenteric, coeliac and mediastinal) were pre-fixed with 0.4 % PFA/PBS *in situ* prior to preparation an full fixation with 4 % PFA/PBS. Afterwards, LN were processed for lightsheet microscopy or cryo sectioning.

### Immunehistochemistry

Pre-fixed LN were washed in PBS, embedded, and snap-frozen in OCT compound. 20 µm cryosections were fixed in 4% PFA/PBS for 15 min, washed in PBS, and blocked in Blocking Solution (10% Chicken Serum, 0.3% Triton X-100 in PBS). Sections were incubated for 1 h with primary antibodies diluted in Dilution Solution (1% BSA, 1% Chicken Serum, 0.3% Triton X-100 in PBS), washed three times in PBS-T (0.1% Tween-20 in PBS), and finally incubated in Alexa-dye-conjugated secondary antibodies. After mounting with Mowiol confocal images were captured.

### Wholemount staining of embryonic and adult tissues

Tissues were fixed in 4 % PFA for 2 h and washed with PBS. Subsequently, samples were permeabilized in 0.5 % Triton X-100 in PBS, blocked in PermBlock solution (3 % BSA, 0.1 % Tween-20 in PBS) and stained with the listed primary antibodies at 4°C. Following three washing steps with PBS-T (0.1 % Tween-20), tissues were incubated in secondary antibodies labeled with Alexa-dyes and mounted with Mowiol.

### β-Galactosidase staining

VEGF-C expression was monitored in embryonic, postnatal and adult tissues by whole-mount staining for β-galactosidase activity. V*egfc*^+/LacZ^ tissues were dissected and fixed in fixative solution (5 mM EGTA, 0.4 % PFA, 2mM MgCl_2_, in 1x PBS) o.n. at 4 °C. The samples were washed with wash buffer (2mM MgCl_2_, 0.02% NP-40, 0.01. % Sodium deoxycholate in 0.1 M Phosphate buffer, pH 7.3) and stained o.n. at 37 °C in staining solution (1 mg/mL X-Gal, 2 mM MgCl_2_, 0.02 % NP-40, 0.01% Sodium deoxycholate, 5 mM Potassium ferricyanide, 5 mM Potassium ferrocyanide in 0.1 M Phosphate buffer, pH 7.3). Thereafter the tissues were washed in wash buffer and recorded using a Nikon D5600 camera (40mm f=2.8 macro lens) or a Nikon Eclipe Ti2-E fluorescence microscope (4x, NA=0.13; 10x, NA=0.3; 20x, NA=0.45; 40x, NA=0.6; 40x, NA=1.15).

### Confocal laser scanning microscopy and image processing

Confocal images were captured using a LSM 880 confocal microscope (Carl Zeiss; 10x, NA=0.45; 20x, NA=0.8; 40x water, NA=1.2; 63x oil, NA=1.4). Microscopy data were recorded and processed with ZEN black (2.3) software (Carl Zeiss). All confocal images represent maximum intensity projections of z-stacks of either single tile or multiple tile scan images. Mosaic tile-scans with 10 % overlap between neighboring z-stacks were stitched in ZEN black (2.3) software.

### Optical tissue clearing

Optical clearing was described previously (Hagerling et al., 2013). Briefly, for visualization, adult lymphnode wholemount stainings were embedded in 2 % low-melting point agarose. After dehydration in methanol (50 %, 70 %, 95 %, 100 %, 100 % methanol, each step min. 1 hour) samples were optically cleared in a benzyl alcohol : benzyl benzoate solution (ratio 1:2) for 4h. Cleared samples were subsequently imaged by lightsheet microscopy.

### Lightsheet microscopy and 3D-image processing

Optically cleared samples were imaged using a LaVision Ultramicroscope II (LaVision BioTec, Bielefeld, Germany). Multi-color Z-stacks were captured with a step size of 1-2μm at different magnifications (Olympus MVPLAPO 2x lense (total magnification 1.26x–12.6x; NA = 0.5). 3D reconstruction and analysis of ultramicroscopy stacks were performed by using the volume rendering software Voreen.

### FACS-sorting

LECs of capillay or collector origin were isolated from adult WT mesenteries (12-14 wks) to subsequently culture the cells and to analyze their lymphatic origin by immunostaining. Simultaneously, SMCs and BECs were isolated from adult WT mesenteries (12-14 wks) to analyze mRNA expression levels of the lymphangiogenic growth factor VEGF-C. Therefore, mesenteries were enzymatically digested in PBS^Ca2+, Mg2+^ supplemented with 100 mg Collagenase A and 1 % Dispase II for 40 min at 37 °C. Single cells were collected and stained for 15 minutes with dye-conjugated primary antibodies diluted in FACS buffer (3 % fetal calf serum in PBS). After antibody staining the cells were analyzed and sorted on a FACSAria IIIu cell sorter (BD Biosciences) using a 85μm nozzle. Cells were gated based on FSC and SSC to exclude cellular debris and doublets, and on DAPI exclusion for viability. CapLEC were sorted on the basis of CD31, Podoplanin and LYVE1 positivity; colLEC via CD31, Podoplanin double positivity and LYVE1 negativity; BECs via CD31 positivity and Podoplanin, LYVE1 double negativity and SMC via CD31, Podoplanin and LYVE1 negativity, but PDGFR-β positivity.

The cells were directly sorted into RLT lysis buffer (Qiagen) and immediately proceeded for total RNA isolation. Data analysis was done using FlowJo software (LLC). To optimize yields two WT mesenteries were pooled for each individual experiment.

### Small scale RNA isolation and cDNA synthesis

Total RNA was isolated from FACS-sorted cells using the spin-column based RNeasy Plus Micro Kit (Qiagen) according to the manufacturer’s instructions. Total RNA was eluted in RNase-free H_2_O and concentration and quality were determined using a 2100 Bioanalyzer (Agilent) and the Agilent RNA6000 Pico Kit and reagents. First-strand cDNA was synthesized from equalized amounts of template RNA using the SuperScript IV Reverse Transcriptase, primed by random hexamers (Life Technologies).

### Preparation of whole tissue RNA lysates for determination of VEGF-C and –D expression levels

Tissue pieces were prepared from the mesentery (excluding the mesenteric LN) of WT and *Vegfc*^+/LacZ^ animals to analyze remaining VEGF-C expression in heterozygously deleted animals and potential compensatory upregulation of VEGF-D. The samples were mechanically dissociated using the Precellys Evolution HP tissue homogenizer (bertin technologies) and the Precellys 2mL tissue homogenizing mixed beads kit (CKMix, Precellys). The dissociation program was run at 6.500 rpm in two repetitive cycles. For subsequent isloation of total RNA, the tissue was directly dissociated in phenol/guanidinium-isothiocyanate buffer (PEQLAB Biotechnologie GmbH).

### Large scale RNA isolation and cDNA synthesis

Total RNA was isolated from tissue lysates in phenol/guanidinium-isothiocyanate buffer (PEQLAB Biotechnologie GmbH) supplemented with 70 µg glycogen (Serva) according to the manufacturer’s recommendations. Total RNA was eluted in RNase-free H_2_O and RNA concentration was spectrophotometrically determinded using the Thermo Scientific™ NanoDrop 2000. First-strand cDNA was synthesized from 2 µg template RNA using SuperScript IV Reverse Transcriptase, primed by random hexamers (Life Technologies).

### Quantitative real-time PCR (qPCR) using TaqMan® Assays

Transcript levels were determined by quantitative real-time PCR (qPCR) using TaqMan® gene expression assays (Applied Biosystems) on a 7300 fast RT-PCR System (Applied Biosystems) with pre-designed exon-spanning reaction mixes (ThermoFisher Scientific). The data was analyzed using the comparative CT method (ΔΔCT) for calculating relative quantitation of gene expression.

### Isolation and culture of murine lymphatic endothelial cells

Primary msn lymphatic endothelial cells (LECs) were isolated from WT mice by FACS sorting. For cultivation of LECs preselected batches of ECGS (BT-203, Alfa Aesar) that favor the expansion and maintenance of lymphatic endothelium were used. In brief, LECs were maintained in Dulbecco’s Modified Eagle’s Medium (DMEM) supplemented with 20 % FCS, 2 mM L-glutamine, 100 μg/mL penicillin/streptomycin, 1 % sodium pyruvate, 0,1 mM non-essential amino acids, 20 μg/mL ECGS, 50 μg/mL Heparin, 50 μM β-Mercaptoethanol and cultured on 0.4 % gelatine coated petri dishes or multi-well plates at 37 °C, 10 % CO_2_ and 95 – 100 % humidity. LECs between passage 1-2 were used for immunofluorescence staining.

### Immunofluorescence staining of LECs

Primary LECs were seeded onto 0.4 % gelatine-coated glass coverslips and grown to intermediate confluence. LECs were fixed in 4 % PFA/PBS for 15 min at RT, washed twice in PBS and permeabilized in 0.5 % TritonX-100 in PBS for 30 min at RT. After blocking in PermBlock (3 % BSA, 0.1 % Tween-20 in PBS) for 1h at RT, the cells were incubated with primary antibodies diluted in PermBlock solution o.n. at 4 °C, washed three times in PBS-T (0.1 % Tween-20 in PBS) and finally incubated in Alexa-dye conjugated secondary antibodies (Life Technologies). After mounting with Mowiol (Calbiochem), confocal images were captured using a Zeiss LSM 880 (20x, NA=0.8; 40x water, NA=1.2; 63x oil, NA=1.4) confocal microscope.

### Single-cell RNA-sequencing

To compare the expression level of Vegfc and other genes of interest among different murine cell types, we obtained single cell RNA sequencing data reported by González-Loyola et al. (2021) (Gonzalez-Loyola et al., 2021). These cells were obtained by dissociating whole mesenteries excluding mesenteric lymph nodes of 3 wild type mice, followed by sorting of live CD45^−^ CD31^+^ PDPN^+^ cells. Single cells were subjected to scRNAseq using the 10x Genomics droplet encapsulation technology. The initial pre-processing steps of the data are described in the Methods section of (Gonzalez-Loyola et al., 2021). The cells presented in Fig. S14E of González-Loyola et al. were subset to keep the normalized counts of wild type cells annotated as blood endothelial cells (BECs), fibroblasts, capillary and lymphangion lymphatic endothelial cells (LECs), mesothelial cells, pericytes and smooth muscle cells (SMCs). To compare the level of Vegfc between cells annotated as BECs 3 and SMCs, we filtered the BECs 3 cells to only retain the ones that had normalized counts>0. Violin plots of showing the distribution of normalized counts per gene within each cell type were generated using the Seurat package (v4.0.3 (Hao et al., 2021) in R (v4.0.3).

### Analysis of publicly available scRNA-Seq data

We examined the presence of LYVE1+ macrophages in two publicly available scRNA-Seq data sets. We obtained the raw counts matrices of murine whole mesenteric cells from the Gene Expression Omnibus (GEO), accession number GSE102665 (Koga et al., 2018), and of murine omentum macrophages from the functional genomics data collection (ArrayExpress), accession number E-MTAB-8593 (Etzerodt et al., 2020). We re-analyzed both data sets similarly to Zhang et al, 2021, using the Seurat package (v4.3.0) for R (v4.2.2). The raw counts were converted to ln (normalized counts +1). Dimensions were reduced using principal component analysis, followed by Uniform Manifold Approximation and Projection (UMAP) on the 10 first principal components, 30 neighbors and the cosine distance metric. Cells were clustered using a shared nearest neighbor (SNN) modularity optimization (Louvain) based clustering algorithm on the 10 first principal components, with 20 nearest neighbors and a resolution parameter of 0.2.

## Quantification and statistical analysis

### Image processing and analysis in fiji

Confocal single and multi tile-scans were processed in fiji. If necessary, adjustments to brightness, contrast and intensity were equally accomplished for individual channels and compared data sets. Quantification of lymphatic vessel parameters (vessel area, branching, diameter, number of sprouts, number of valves) was done in fiji using maximum intensity projection images. The cell counting plugin was used to assess the number of branchpoints and valves. Details on image analysis can be found in the according figure legends.

### Statistical analysis

Statistical analyses were perform with GraphPad Prism 8 software. Significance was analyzed using two tailed unpaired Student’s t-tests with Welch’s correction for assumption of unequal SD. The significance threshold was set at 0.05. P-values are indicated by asterisks: *, p ≤ 0.05; **, p < 0.01; ***, p < 0.001 and ****, p<0.0001. Results are shown as mean ± SD. As indicated in the respective figure legends, all representative data were confirmed in three or more independent experiments.

## Additional resources

### Key resources table

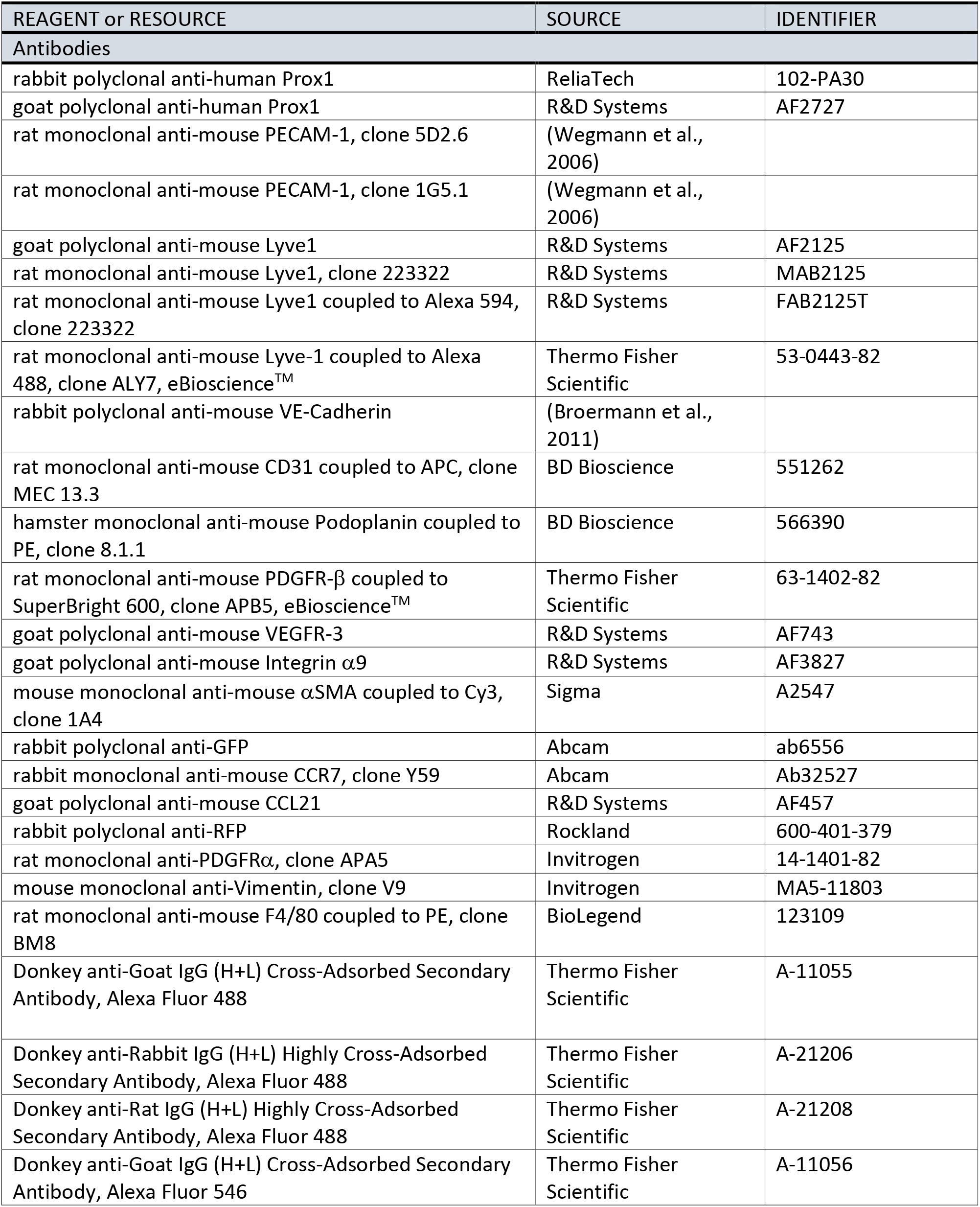
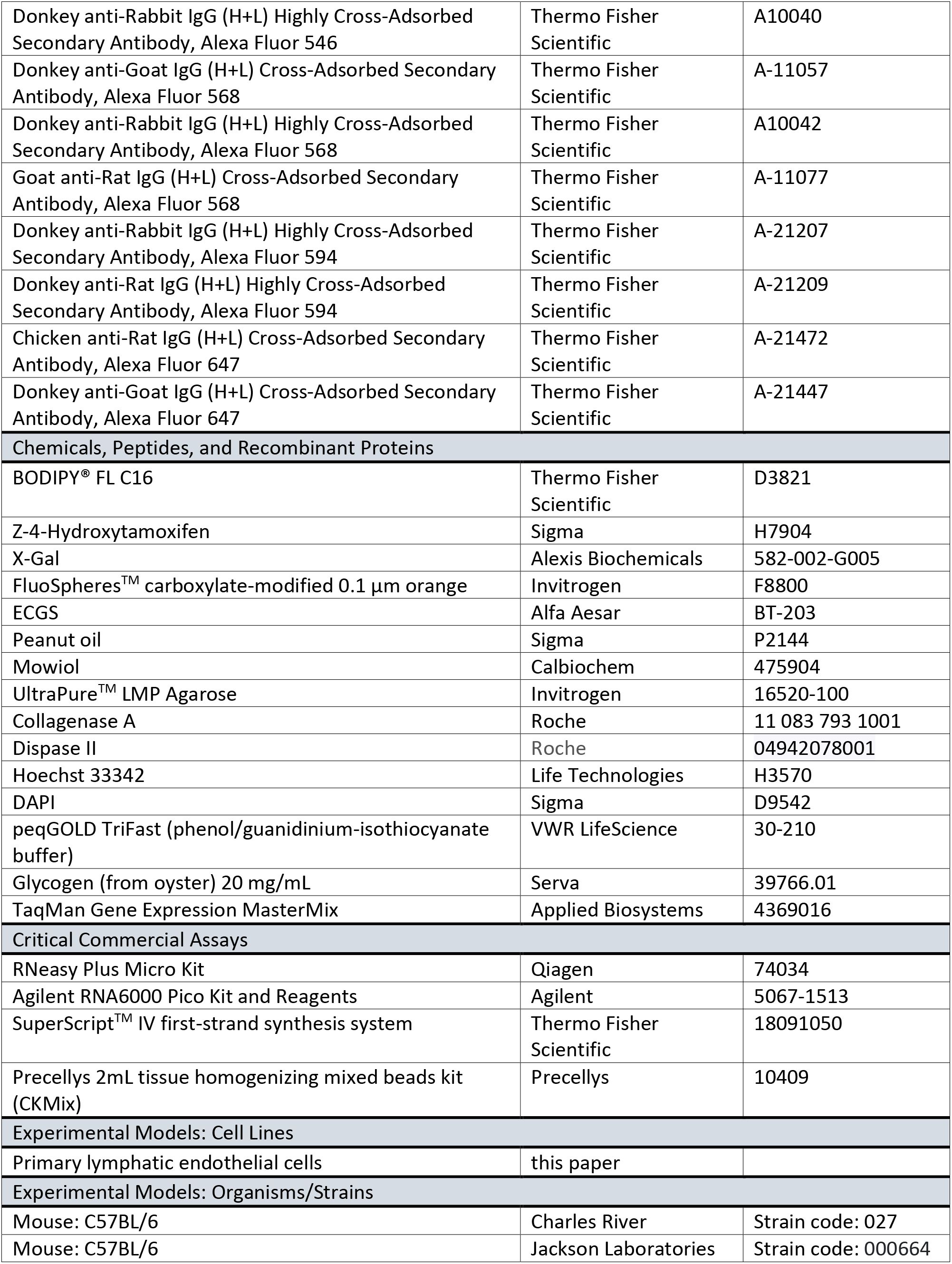
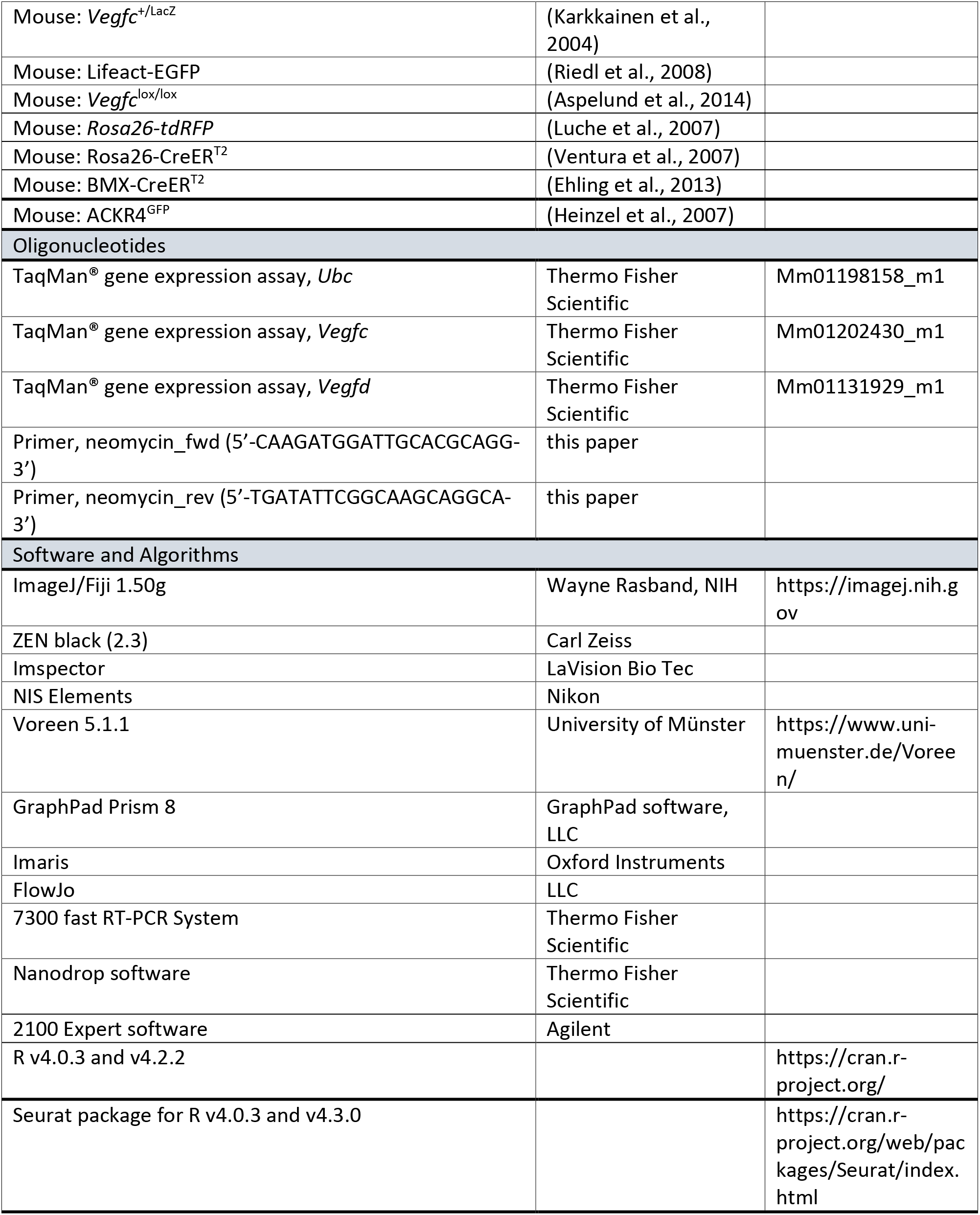

