## Supplemental Figures and Legends for "Specialized mesenteric lymphatic capillaries by-pass the mesenteric lymph node chain to transport peritoneal antigens directly into mediastinal lymph nodes"

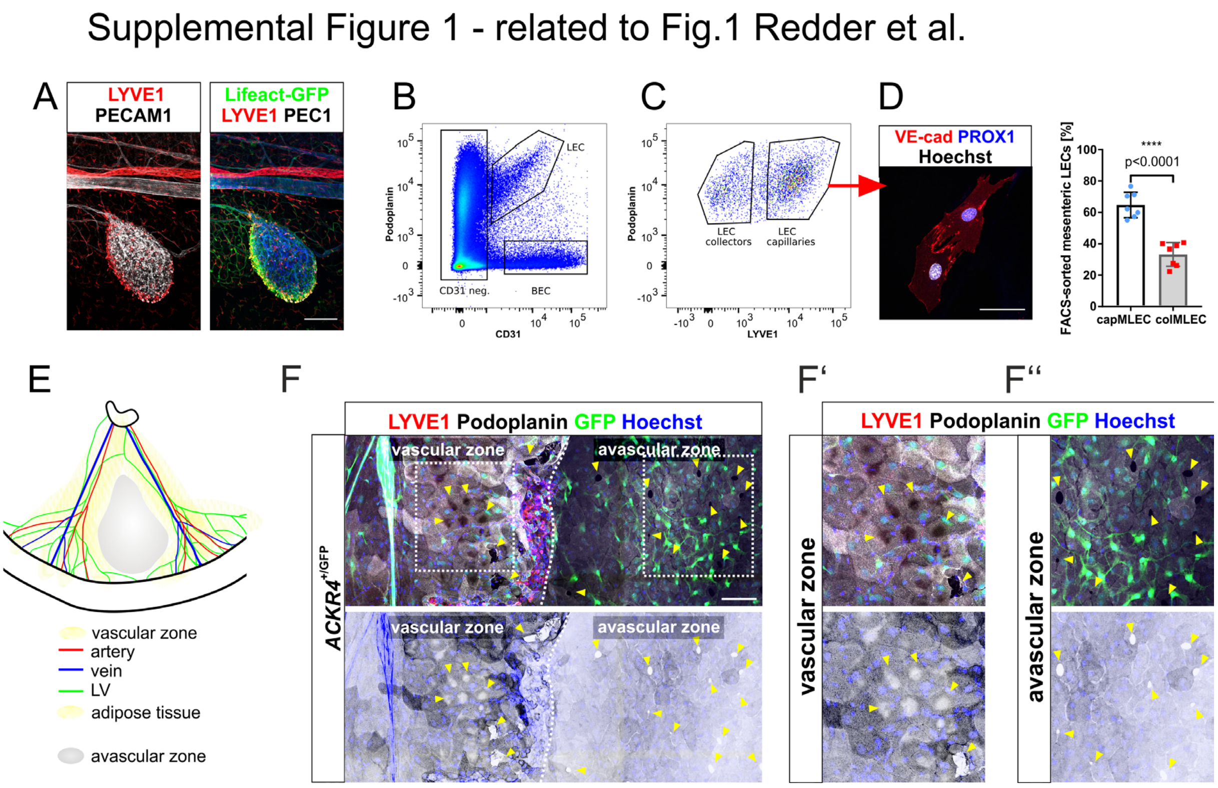


Figure S1 related to Figure 1

**Characterization of mesenteric lymphatic capillaries and their association with FALCs and stomata**

**A** Magnification of a mesenteric FALC (small box in Fig. 1A) showing rich blood vascularization, while the next lymph capillary (Lyve1+) is approx. 200µm apart. Scale bar = 200 µm.

**B-C** Heat-map including the gates for FACS of mesenteric LECs and BECs based on co-expression of PECAM1 (CD31) and podoplanin (B), capillary mesenteric LECs (capMLECs) were distinguished from collector LECs (colMLECs) based on LYVE1 expression (C).

**D** Sorted capMLECs expressed VE-cadherin and PROX1 in culture, demonstrating their LEC identity. Scale bar = 50 µm. The bar diagram indicates the relative abundance of capMLECs and colMLECs in sorted mesenteric preparations. Data represent mean (*n*=7) ± s.d. Statistical significance $****P \leq 0.0001$. Two-tailed unpaired Student's *t*-test with Welch's correction.

**E** Schematic representation of a mesenteric segment indicating the vascular zone (vascular bundle and adipose tissue) and large intermittent avascular zones.

**F-F''** MIPs (multi-tile z-stack) of a wholemount immunostained adult *ACKR4*$^{+/GFP}$ mesentery depicting the presence of stomata (yellow arrows) in the vascular and avascular zone of the mesentery. Mesothelial cells were stained with podoplanin (grey scale), the white dotted line marks the border of the fat-covered vascular zone (with rare ACKR4+ cells, green) and the avascular zone (rich in ACKR4+ cells, green). In the bottom panel the LUT for podoplanin (grey) is inverted to highlight the position of stomata (distinct white spaces). The boxed areas in E are shown magnified in E' and E'' (yellow arrowheads denote stomata in both zones). Stained are the antigens in indicated color. Scale bar = 200µm.

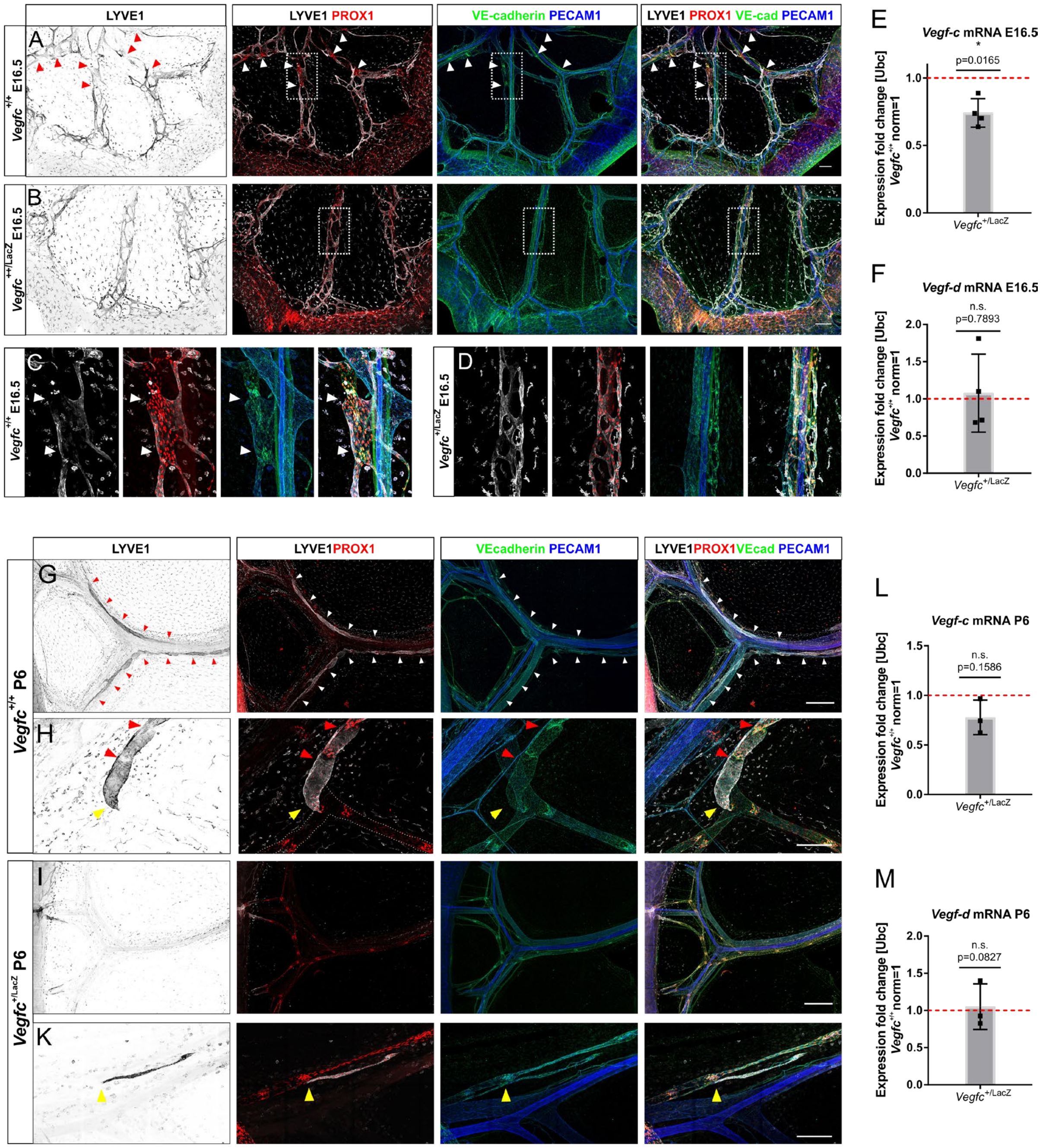


Figure S2 related to Figure 2

**Development of the fetal mesenteric lymphatic vasculature is impaired in mice heterozygous for *Vegfc*.**

**A - D** MIPs (multi-tile z-stack) of wholemount immunostained fetal mesentery of E16.5 control (*n*=5) and *Vegfc*$^{+/LacZ}$ (*n*=4) littermates. Antigens are depicted in color above panels (VE-cad=VE-cadherin). Arrowheads in A and C indicate areas rich in PROX1-high- and LYVE1-low expressing LECs, which indicate the onset of valve formation and hence collecting vessel maturation. The boxed areas in A and B are magnified in C and D. Scale bars = 200 µm.

**G - K** Multi-tile MIPs of mesenteric wholemount immunostainings of P6 control (*n*=3) and *Vegfc*$^{+/LacZ}$ (*n*=2) littermates. Antigens are depicted in color above the panels in the top row (VE-cad=VE-cadherin). Arrowheads in (G) follow the expanding capMLV. **H** Elongated capMLVs are still attached to colMLVs (yellow arrowhead) and display transient intraluminal valves (red arrowheads). **I, K** In *Vegfc*$^{+/LacZ}$

mesenteries only rudimentary capMLV are observed (yellow arrowhead in K). Scale bars = 500µm (G, H); 200 µm (I, K).

**E-F, L-M** qRT-PCR-based transcriptional analysis of the lymphangiogenic growth factors VEGF-C (E, L) and VEGF-D (F, M) in E16.5 (E,F) and P6 (L,M) control and *Vegfc*$^{+/LacZ}$ mesenteries. Individual samples were measured in triplicate. Depicted is expression of target genes relative to the control transcript *Ubc* ($\Delta\Delta C_t$). *Vegfc*$^{+/+}$ expression levels were normalized to 1 (red dotted line). Data represent the mean ± s.d (*n* = 4 E-F, n = 3 L, M). Statistical significance is noted above each graph. Two-tailed unpaired Student's *t*-test with Welch’s correction.

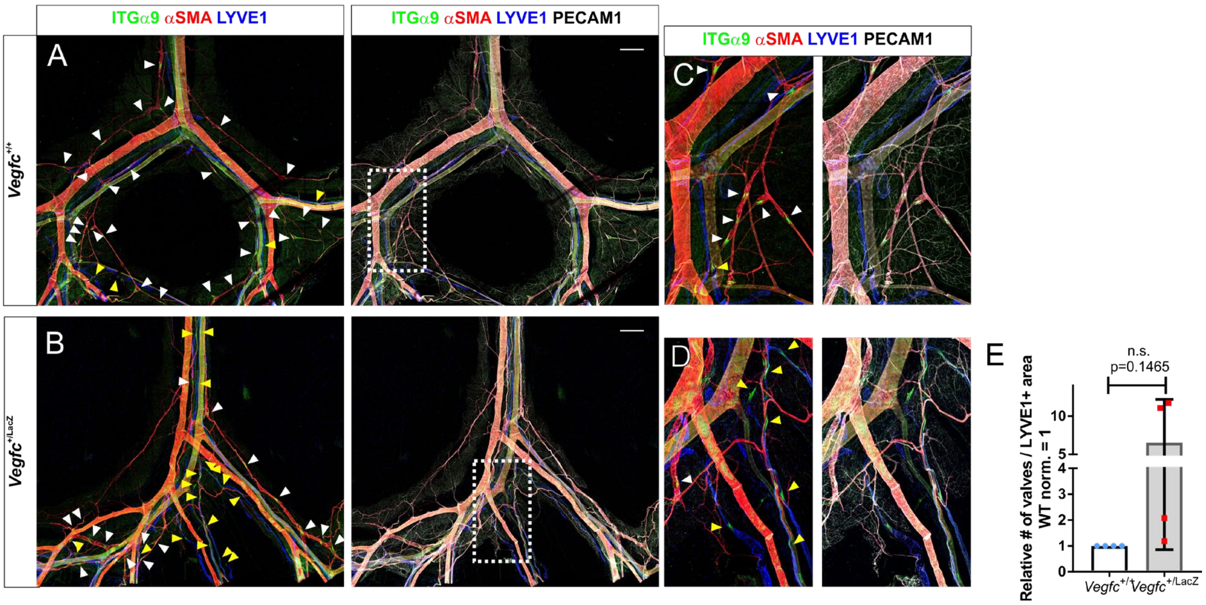


Figure S3 related to Figure 3

**Hypomorphic architecture of lymphatic capillaries and increased frequency of intraluminal valves in *Vegfc*$^{+/LacZ}$ mesenteries indicate a developmental delay that results in an increased frequency of pre-collector-type vessels.**

**A - D** Multi-tile MIPs of wholemount stained mesenteries prepared from adult control (A, C) or *Vegfc*$^{+/LacZ}$ mice (B, D). (C) and (D) show magnifications of the areas outlined (white dashed rectangle) in (A) and (B). High ITGα9 expression identified lymphatic valves. White arrows, valves in lymphatic collectors; yellow arrows, valves in lymphatic capillaries (A-D). Scale bar = 500 µm.

**E** Analysis of the valve frequency in capMLVs normalized to the LYVE1-positive area. Data represent the mean ± s.d ($n$ = 4). WT values were normalized to 1**.** Statistical significance is noted on top. Two-tailed unpaired Student's $t$-test with Welch's correction.

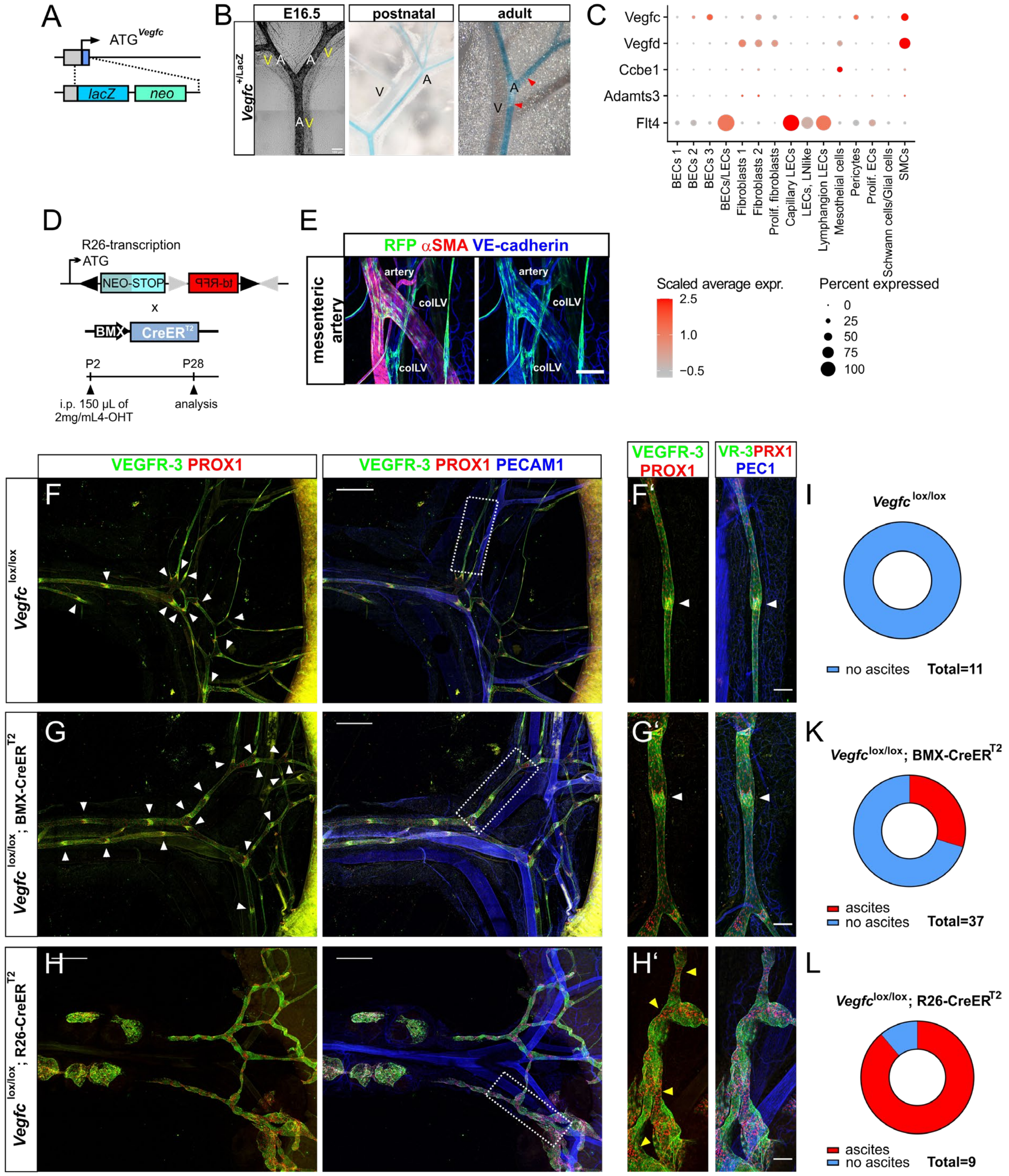

Figure S4 related to Figure 4

**Differential sensitivity of colMLVs and capMLVs to reduced VEGF-C expression. Partial loss of arterially expressed VEGF-C results in absence of capMLVs, while colMLVs decay only after global postnatal VEGF-C deletion.**

**A** Schematic of the genetically modified VEGF-C locus employed during visualization of mesenteric VEGF-C expression based on LacZ/β-Gal staining.

**B** VEGF-C expression in mesenteric arteries identified by β-Gal staining in preparations isolated at the indicated stages from *Vegfc*$^{+/LacZ}$ mice (A = artery, V = vein). Note E16.5 representation is in greyscale. Scale bar = 100 µm. Red arrowheads point to circumferential expression pattern in adult samples.

**C** Dot plot representing the expression of selected genes related to VEGF-C signalling in mesenteric cell subsets of adult mice. The color code indicates the scaled average expression level, while the dot size indicates the percentage of cells expressing the given gene in each cluster.

**D** Schematic depiction of the *R26-tdRFP* reporter allele for Cre activity. Cre-mediated recombination of incompatible loxP and loxP2272 sites result in inversion of the tandem-dimer red fluorescent protein sequence and excision of the neo/stop codon cassette permitting expression of the non-toxic tdRFP (two modified DsRed subunits covalently linked). Below: Regime for postnatal 4-OHT administration to trigger activation of Cre (BMX-CreER$^{T2}$)-recombinase in arterial ECs (BMX-CreER$^{T2}$) at early postnatal development.

**E** BMX-CreER$^{T2}$-mediated expression of tdRFP in mesenteric arterial endothelial cells of R26-tdRFP; BMX- CreER$^{T2}$ mesenteries. MIPs generated from single-tile scans show incomplete recombination in arterial ECs. Stained antigens are indicated in color above each panel. Scale bar = 100 µm.

**F – H'** MIPs (overview tile-scans) of P28 *Vegfc*$^{lox/lox}$ (*n*=4), *Vegfc*$^{lox/lox}$; BMX-CreER$^{T2}$ (*n*=4) and *Vegfc*$^{lox/lox}$; R26-CreERT$^{2}$ (*n*=5) mesentery wholemount preparations immuno-stained for the indicated antigens (VR-3=VEGFR-3, PRX1=PROX1, PEC1=PECAM1). The white boxed areas in (F - H) are magnified in (F' – H'). White arrowheads in (F, F' – G, G') point to intraluminal valves characterized by high PROX1 and VEGFR-3 expression and leaflet formation. Yellow arrowheads in (H') emphasize areas of high PROX1 expression, which lack intact leaflet formation in colMLVs following ubiquitous postnatal *Vegfc* deletion. The overviews in (H) denote LEC-evaginations in colMLV and unspecific accumulation of PROX1+ LECs. Scale bar = 500 µm (F-H); 100 µm (F'-H').

**I-L** Occurrence of ascites in 4-OHT treated control, *Vegfc*$^{lox/lox}$; BMX-CreER$^{T2}$ and *Vegfc*$^{lox/lox}$; R26-CreER$^{T2}$ mice at P28.

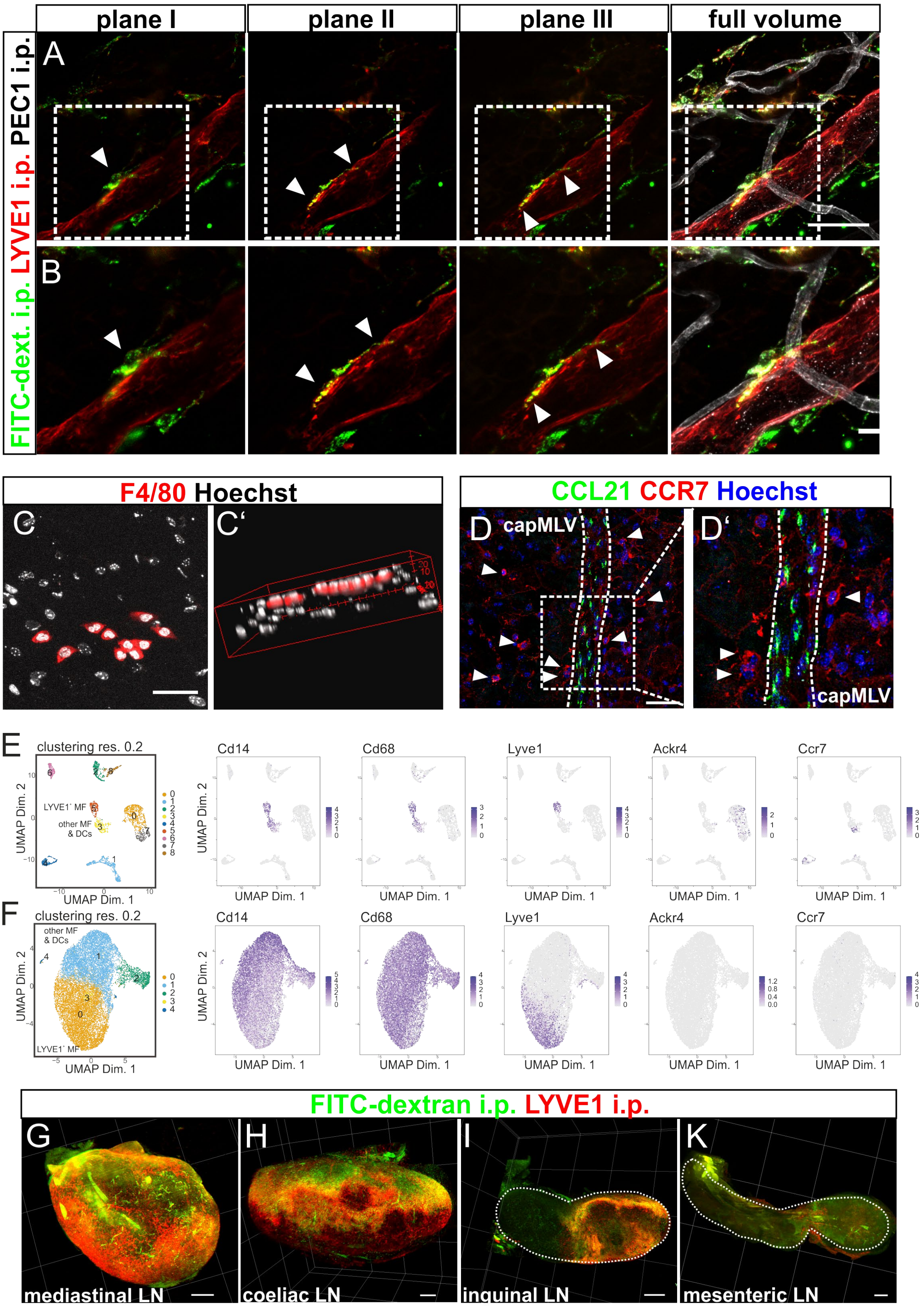


Figure S5 related to Figure 5

**Intravasation of LYVE1-positive phagocytes into capMLVs.**

**A-B** Immunostained mesenteric wholemount preparation 45 min after intraperitoneal FITC-dextran injection identifies phagocytes entering a capMLV (white arrowheads). Antigens are shown in color above each panel. Shown are single z planes of an image stack and MIPs of the full volume. The boxed areas in A are magnified in B. Scale bar = 50µm.

**C-C'** MIP of an overview tile-scan of the avascular area from a P28 WT (*n*=6) mesenteric wholemount preparation immuno-stained for the macrophage marker F4/80 (C). F4/80+ peritoneal fluid macrophages are distinguishable from tissue resident mesenteric phagocytes by F4/80 immunoreactivity, oval morphology and localization atop the mesothelium (C', 3D volume). Scale bar = 50µm.

**D-D'** MIPs (overview tile-scan) of a P28 WT (*n*=3) mesenteric wholemount preparation immuno-stained for the indicated antigens. The white boxed area in (D) is magnified in (D'). The white dotted line outlines a CCL21 positive capMLV. White arrowheads point to CCR7+ phagocytes. Scale bar = 50µm.

**E** UMAP plot displaying reanalysis of scRNA-seq of whole mesentery cells (GSE102665). Expression levels of Cd14, Cd68, Lyve1, Ackr4 and Ccr7 overlaying the UMAP plots are shown on the right.

**F** UMAP plot displaying reanalysis of scRNA-seq of omental macrophages (E-MTAB-8593). Expression levels of Cd14, Cd68, Lyve1, Ackr4 and Ccr7 overlaying the UMAP plots are shown on the right.

**G-K** Digital volume rendering of mediastinal (G), coeliac (H), inguinal (I) and mesenteric (K) lymph nodes 90 min after FITC-dextran i.p. injection (n=6). FITC-dextran and LYVE1 antibodies were detected within mediastinal, coeliac and inguinal LN but not mesenteric LN. Scale bars = 660 µm.

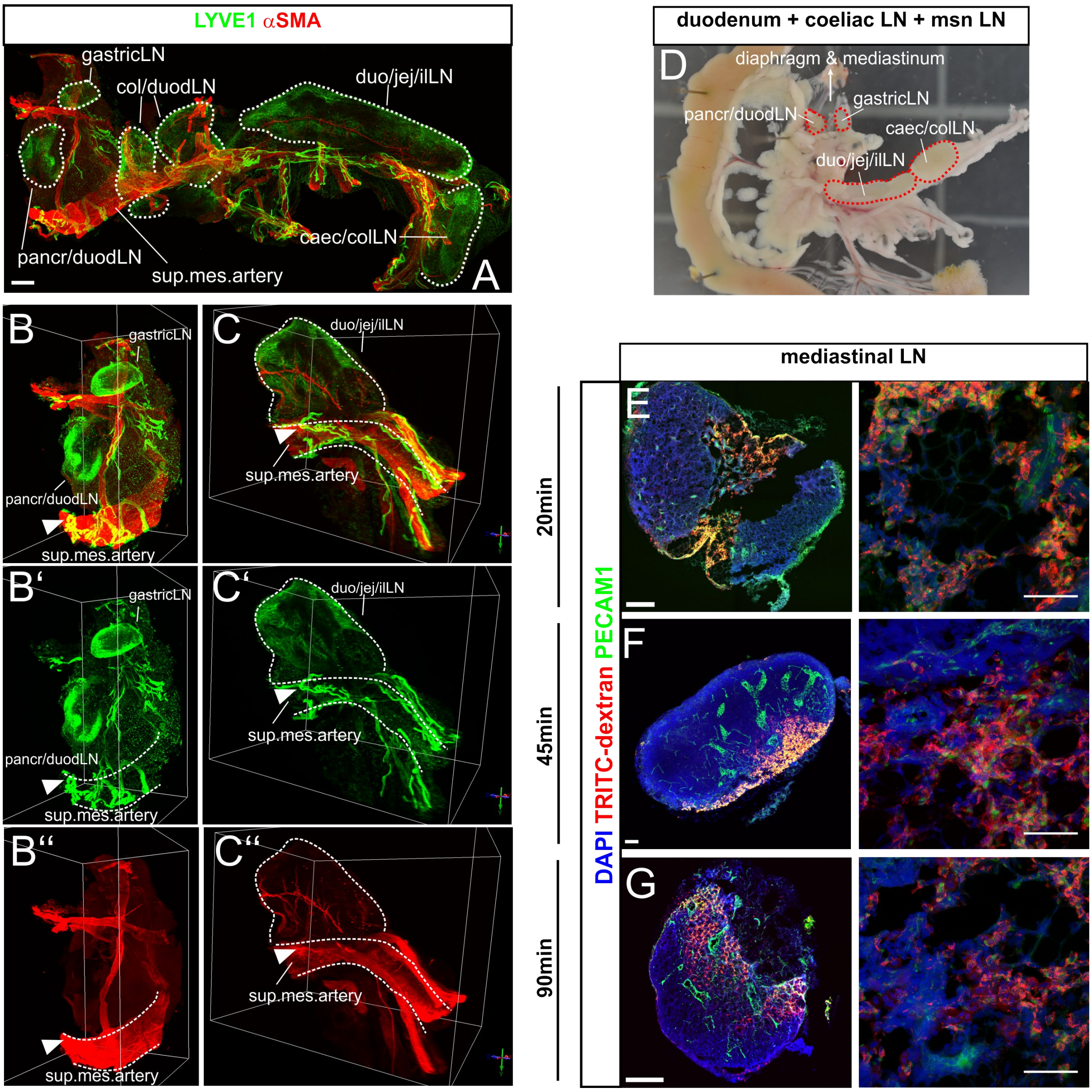


Figure S6 related to Figure 6

**Identification of mediastinal lymph nodes draining mesenteric lymphatic capillaries.**

**A – C''** Digital volume renderings of light sheet microscopic image stacks from full wholemount preparations immunostained for LYVE1 and αSMA. Two selected volume aspects are shown in B – B'' (coeliac LN = gastric and pancreatoduodenal node) and C – C'' (mesenteric LN = duodenal, jejunal and ileal nodes). White dotted lines outline the superior mesenteric arteries and follow the shape of the indicated LN. White arrowheads in B-C'' indicate mesenteric capLVs that follow the superior mesenteric artery towards the abdominal aorta. The data represents four individual preparations.

**D** Stereomicroscopic view of a full wholemount preparation including a large part of the duodenum, the mesentery and the indicated fat embedded lymph nodes (LN). We refer to the gastric and pancreatoduodenal node collectively as coeliac nodes.

**E – G** MIPs of immune-labelled 20 µm cryosections of mediastinal LN after intramesenteric injection of TRITC dextran and i.p. injection of PECAM1 antibodies. Times of analysis after TRITC dextran injection are indicated. Scale bar = 500 µm, 50 µm (right panels).

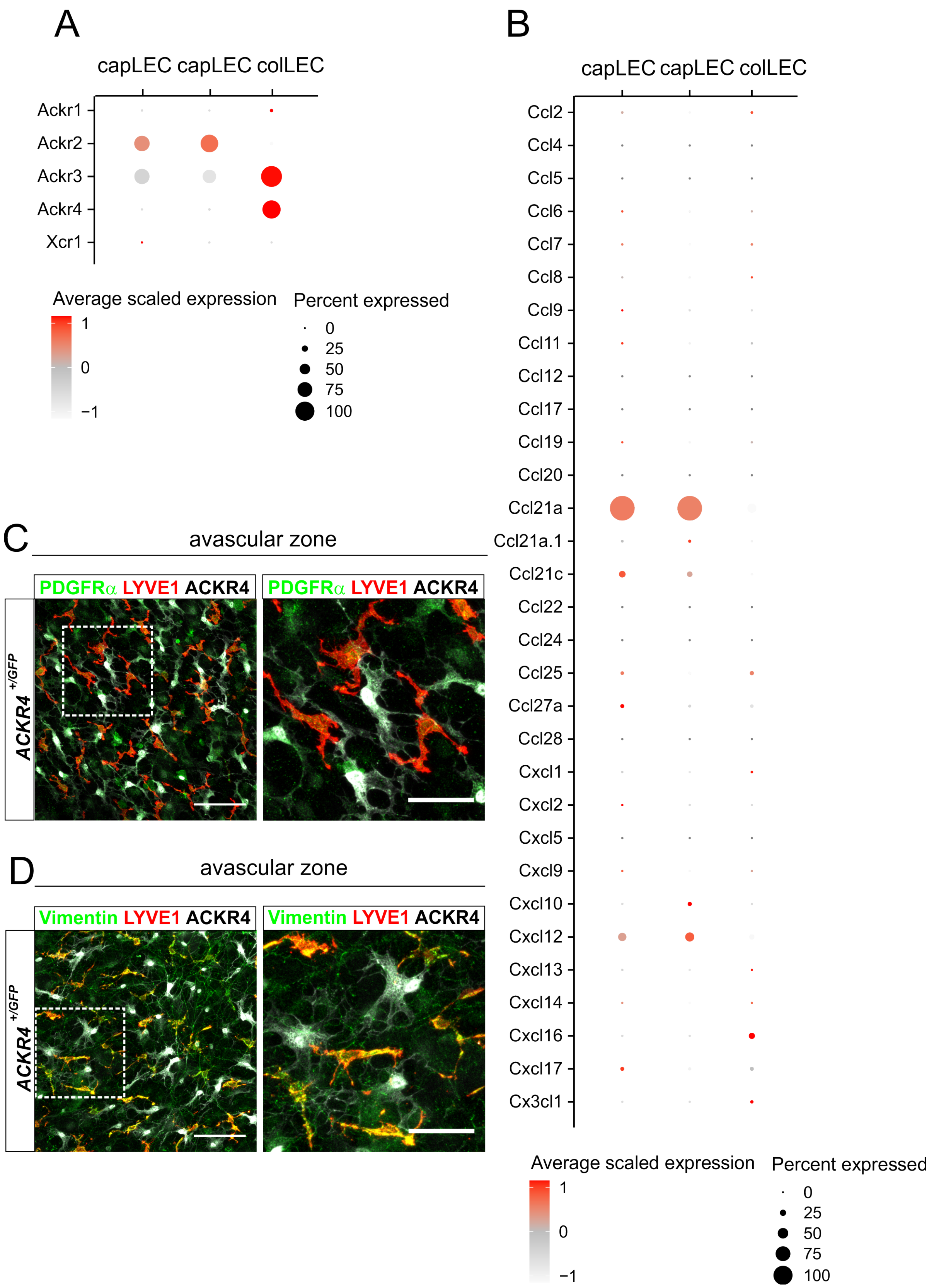


Figure S7 related to Figure 7

**Chemokine and chemokine receptor expression in capMLEC and colMLEC**

**A - B** Dot plot of selected genes encoding atypical chemokine receptors and chemokines in two mesenteric capMLEC and one colMLEC subsets of adult WT mice. The color code indicates the scaled average expression level, while the dot size indicates the percentage of cells expressing the given gene in each cluster.

**C - D** Characterization of resident cells located in the avascular regions of the mesentery. Shown are MIPs of overview tile-scans of WT (*n*=3) mesenteric wholemount preparations immuno-stained for the indicated antigens. The white boxed areas in (C-D are magnified in the right panels. Scale bar = 100µm (C, D), 50µm (magnifications).
